# Single anticodon-edited tRNA therapy targeting highly prevalent Arg>Ter premature termination codons causing inherited retinal diseases

**DOI:** 10.64898/2026.07.21.737205

**Authors:** Asodu Sandeep Sarma, Alaa Saleh, Jonathan Eintracht, Hala Kamal, Samer Khetab, Manar Salameh, Chen Matsevich, Alexey Obolensky, Eyal Banin, Dror Sharon

## Abstract

Nonsense variants cause 18% of inherited retinal diseases (IRDs), yet current therapies require variant-specific development, leaving most patients untreated. Here, we combined a large-scale genetic analysis literature survey of >37,500 IRD patients with anticodon-edited (ACE)-tRNA engineering to create a single, gene-agnostic therapy targeting Arg>Ter nonsense variants which are the most prevalent subclass (35%) of premature stop codons (PTCs). We developed an optimized ACE-tRNA (V3) that achieved up to 86% readthrough across 13 clinically relevant variants, restored native PRCD localization in the arRP-causing p.R22* mutant, and demonstrated activity in photoreceptor-like cells. To enable translation, we established an AAV2/7m8 production platform (1*10¹²–1*10¹³ GC/mL) and defined 1*10 GC/eye as the safe dose in mice. This patient genetics-guided strategy positions ACE-tRNA_V3 as a promising candidate for preclinical development, offering a precision medicine approach that targets the most common nonsense variant class with a single therapeutic molecule.

## Introduction

Inherited retinal diseases (IRDs) are clinically complex and genetically heterogenous rare visual impairment disorders that affect approximately 1 in 1380 individuals and account for a significant proportion of visual impairment and blindness worldwide ^1,2^. IRDs are characterized by progressive degeneration of photoreceptors or other retinal cell types and can manifest at any age ^3^. More than 350 genes have been implicated in their pathogenesis with *ABCA4, USH2A, EYS, RPGR* and *CRB1* being frequently mutated genes in IRD patients ^4,5^. More than 50 different types of IRD phenotypes have been reported in the literature with retinitis pigmentosa being the most common subtype ^6,7^. Premature termination codons (PTCs) account for around 11% of the all-rare disease-causing variants ^8^ and 18% of IRD-causing variants ^4^. PTCs usually activate a highly conserved cellular surveillance mechanism, nonsense-mediated mRNA decay (NMD), resulting in mRNA degradation and loss-of protein function ^9,10^. However, in some instances transcripts with PTC escape NMD and are translated into a truncated protein resulting in either loss or gain of protein function ^11,12^.

In recent years treatment of IRDs has significantly advanced, most notably through the approval of gene augmentation therapy for Leber congenital amaurosis caused by biallelic *RPE65* pathogenic variants ^13–16^. However, gene augmentation therapy is limited by the AAV cargo packing capacity, the elicited immune response and the risk of long-term gene over-expression ^17,18^. Other recent developments include genome editing, RNA editing, antisense oligonucleotides (AONs) and cell-based therapies, each with its own advantages and challenges ^4^. These approaches are largely gene- or variant-specific, meaning that each one requires a separate development pipeline. This creates a very restrictive bottleneck for treating IRDs as the extensive time and cost of *in vitro* and *in vivo* testing, followed by clinical trials and FDA approval protocols, must be navigated individually for each novel therapy. While gene-specific therapies have transformed treatment for some IRD patients, the extreme genetic heterogeneity of these conditions makes personalized development economically unviable for most. A variant class-based strategy that addresses multiple genes simultaneously would represent a paradigm shift.

Anti-codon edited tRNAs (ACE-tRNA) offer a potential alternative for nonsense variants readthrough. ACE-tRNAs are derived from natural tRNAs with an altered or edited anticodon which can base-pair with one of three nonsense codons (UAG, UAA or UGA). ACE-tRNAs are charged with their cognate amino acids and participate in translation elongation at the targeted PTC, which allows the production of full-length wild-type protein from a PTC-containing mRNA ^19^. In recent years, several studies have demonstrated proof-of-concept for ACE-tRNA mediated translational readthrough both in vitro and in vivo ^19–22^, suggesting that this approach can overcome several challenges posed by other readthrough agents such as the risk of global readthrough of natural stop codons ^23^.

Due to current limitations of existing translational readthrough strategies, we explored the potential of ACE-tRNAs to suppress common IRD-causing nonsense variants and restore full-length protein production. Although the ACE-tRNA-based PTC suppression strategy has shown promise in other rare diseases such as cystic fibrosis and others ^20–22^, their application to IRDs has been limited by a lack of systematic analysis identifying the most clinically relevant PTC subclasses^24,25^.

In this study, we initially analyzed published genetic data from over 37,500 IRD patients and identified Arg>Ter nonsense variants as the major disease-causing subclass. Guided by this insight, we generated an optimized ACE-tRNA targeting this subclass and validated its efficacy across 13 most prevalent nonsense variants, culminating in a safe and AAV-deliverable therapy. These findings establish a gene agnostic precision-medicine strategy for the most common class of nonsense variants in IRDs.

## Methods

### Collecting IRD genetic data from the literature

We extended the previously published GRID dataset ^4^ to include a larger number of IRD cases. The dataset was analyzed for frequency of various PTCs.

### Cell culture and transient transfection

HeLa cells were maintained in Dulbecco’s Modified Eagle’s Medium (DMEM) supplemented with 10% Fetal bovine serum (FBS), L-glutamine, and penicillin/streptomycin and grown at 37 C and 5% CO2. Cell lines were routinely tested for mycoplasma contamination. All transfections were carried out using TOCRIS PEI STAR transfection reagent (Cat. No. 7854).

### Plasmids and cloning

The pmCherry-GFP (#86639) and EGFP-N2 plasmids were obtained from Addgene. For the read-through screening of 13 distinct PTC variants, partial cDNA sequences (approximately 150 bp on either side of the PTC) along with the full-length cDNA of *PRCD* were synthesized as g-blocks by IDT. These cDNA fragments were then inserted into the pmCherry-GFP plasmid using the BamHI and KpnI restriction sites. For PRCD localization assay, full-length cDNA was cloned into the EGFP-N2 plasmid using the BamHI and KpnI restriction sites. All three variants of ACE-tRNA^Arg^ were synthesized as g-blocks by IDT and subsequently cloned into two different AAV plasmids, pDS001-mCherry and pDS002-STF **(Supplementary Table 4).** For the experiments conducted in 661W cells both ACE-tRNA^Arg^ and both *FAM161A* wildtype and mutant cDNA fragments were cloned into pDS003-mCherry plasmid. All clones were sanger sequenced and confirmed. All the cDNA sequences used in this study are listed in **Supplementary Table 2.**

### Immunofluorescence and high-resolution fluorescent microscopy

HeLa cells grown on coverslips (70-80% confluence) were transfected using PEI STAR reagent (Cat No. 7854). After 48 hours post-transfection, cells were fixed with 4% PFA, permeabilized with 0.25% Triton X-100, and blocked with 1% BSA. Coverslips were incubated overnight at 4°C with primary antibodies, followed by secondary antibodies for 2 hours at room temperature. Coverslips were mounted with DAPI-containing mounting media (FluroMounter with DAPI Bio SB Cat: BSB0164). Slides were observed under Nikon R1A confocal microscope. All the antibodies used in this study are listed in **supplementary file 6.**

### Incucyte

For Incucyte imaging, HeLa cells were cultured in 24-well plates until they reached 80% confluence. The cells were then transfected with different plasmids using TOCRIS PEI STAR transfection reagent. After 24 hours, the cells were washed with 1x PBS, and fresh media was added. Forty-eight hours post-transfection, the cells were washed again with 1x PBS, fresh media was added, and then imaged using the Incucyte live cell imaging system (Incucyte® SX5 Live-Cell Analysis System, Sartorius). Fluorescence intensity was measured using the built-in Incucyte software.

### FACS

For FACS analysis, HeLa cells were cultured in 24-well plates until they reached 80% confluence. The cells were then transfected with different plasmids using TOCRIS PEI STAR transfection reagent. After 48 hours post-transfection cells were washed with 1x PBS, dissociated with 25% trypsin-EDTA (Merck: T4049), pelleted at 1,000 rpm for 5 min, resuspended with PBS with calcium and magnesium (Merck: D8537). Cells were then sorted using the cytoflex cell sorter, and data were analyzed by using FlowJo.

### Western blot

Cells were harvested 48 hours post-transfection and lysed in RIPA buffer containing protease inhibitors (Merck: R0278). Protein lysates were quantified, separated by SDS-PAGE, and transferred onto PVDF membranes. Membranes were blocked with blocking buffer (Biorad EveryBlot: 12010020) for 5 min, incubated overnight at 4°C with primary antibodies, followed by HRP-conjugated secondary antibodies for 1 hour. Protein bands were visualized using ECL substrate (Biorad:1705060S) and imaged on a Biorad chemiluminescence system.

### MTT cell viability assay

Cells were seeded in 24-well plates and treated as indicated. After 48 hours post-transfection, MTT solution (0.5 mg/mL) was added to each well and incubated for 3–4 hours at 37°C. The formazan crystals were dissolved in DMSO, and absorbance was measured at 570 nm using a microplate reader. Cell viability was calculated relative to untreated controls.

### AAV2/7m8 vector production

For low-cis triple transfection method, the cis plasmid concentrations were reduced to 2%, 10%, and 15%. AAV vectors were produced as previously described ^26^. Briefly, 72h following transfection, viral particles were harvested from supernatant and cell lysate, treated with DNase (11284.932.001, Roche, Switzerland) and purified on an iodixanol density gradient (OptiPrep Density Gradient Medium, #D1556, Sigma-Aldrich, USA). Further desalting, purification, and concentration of the vectors was performed on Amicon Ultra-15 (Millipore, USA) to resuspend the vectors in PBS + 0.001% Pluronic F-68 (#P5556, Sigma-Aldrich, USA). Concentration of the vector preparation was determined by quantitative PCR^56^ and expressed as genome copies (GC) per millilitre (ml).

### Subretinal injections in mice

The study followed the Association for Research in Vision and Ophthalmology (ARVO) Statement for the Use of Animals in Ophthalmic and Vision Research and was approved by the Hebrew University-Hadassah institutional animal care and use committee (IACUC). Subretinal injections were performed as previously described ^26^. Briefly, mice were anesthetized with an intraperitoneal injection of ketamine (85 mg/kg) and xylazine (15 mg/kg). Under direct visualization with a dissecting microscope (Carl Zeiss, Oberkochen, Germany) the head of the mouse was stabilized using a mouse head holder (921-E, David Kopf Instruments, Tujunga, CA, USA). A sclerotomy was made just posterior to the limbus using a 30G needle. A 34G blunt needle connected to a Hamilton syringe was inserted through the sclerotomy and advanced to the subretinal space. A total of 1 µL of viral vector suspension at three different titers was injected slowly to generate a subretinal bleb. Successful bleb formation was confirmed visually. At 5 weeks post injection, retinal structure was studied in vivo by optical coherent tomography (Heidelberg-SPECTRALIS OCT) and fundus autofluorescence (FAF) imaging. The procedures were performed in anesthetized mice with dilated pupils as described previously ^27^.

### Immunohistochemistry (IHC)

Eyes were enucleated, fixed in Davidson solution for 8h at 4°C, and transferred to 70% ethanol overnight at 4°C. Eyes were then processed, embedded in Paraplast and serially sectioned (5-mm thickness). For IHC, deparaffinized sections were incubated in a decloaking chamber (Biocare Medical, Pacheco, CA, USA) with 10 mM citrate buffer (ImmunoRetriever 20X with citrate pH 6.62, BioSb) at 125°C; blocked with PBS solution containing 1% bovine serum albumin, 0.1% Triton X-100, and subsequently incubated overnight with mouse anti-mCherry primary antibody (Sigma-Aldrich AB356482). After washing in PBS, specimens were incubated for 1h at room temperature with donkey anti-rabbit secondary antibody (1:250, Cy3, 711-165-152, Jackson ImmunoResearch Laboratories). All the antibodies used in this study are listed in **Supplementary table 6**

### Induced pluripotent stem cell reprogramming, culture and characterisation

Fibroblasts were reprogrammed with the Stemgent ® StemRNA ™ 3 rd Gen Reprogramming Kit (Reprocell, USA, cat#00-0076) according to manufacturer’s recommendations, except that RNA transfections using Lipofectamine ® RNAiMAX ™ Transfection Reagent (Thermo Fisher Scientific, USA, cat#13778030) were performed every other day. Cells were plated on wells coated with Bio-laminin 521 LN (BioLamina, Sweden, cat#LN521-05) and cultured in mTesR Plus (STEMCELL Technologies, Canada, cat#100-0276). Reprogrammed colonies emerged between 10-20 days following transfection and were excised from the culture and maintained in mTesR Plus. Each isolated colony was considered a new clone and maintained accordingly in mTesR Plus and passaged every 3-5 days. Cells were used for differentiation from passage 12 and onwards. For characterization, iPSCs underwent immunocytochemistry analysis for pluripotency marker OCT4 and SSEA4 expression and for ectoderm, endoderm and mesoderm marker expression following spontaneous differentiation of embryoid bodies. Pluripotency was also confirmed by qRT-PCR analysis of pluripotency markers.

### Retinal Organoids (ROs) Differentiation

Cells were grown to 60-80% confluency in 6-well plates (Greiner, Austria, cat#657160). At this point, cells were maintained in E6 ™ media (ThermoFisher Scientific, USA, cat#A1516401) for 2 days. At day 2, cells were transitioned to an induction media comprised of E6 ™ media, 1X N2 supplement (ThermoFisher Scientific, USA, cat#17502048) and 5mM Nicotinamide (Sigma-Aldrich, USA, cat#N0636). From approximately 4 -8 weeks of differentiation, rudimentary neural retina rosettes resembling early optic vesicles emerged in the monolayer. They were excised and cultured in individual low-adherence 96-well plates (Greiner Bio-One, Austria, cat#650970) (Facellitate, Germany, cat#F202003). From day 28 to 35, early optic vesicles were cultured in DMEM/F12 (ThermoFisher Scientific, USA, cat# 31331028), 2% B27 supplement (50X) (ThermoFisher Scientific, USA, cat#12587010), 1X NEAA, 1% Glutamax (ThermoFisher Scientific, USA, cat# 35050061), 1% P/S and 10ng/µL FGF2 (Peprotech, USA, cat#100-18b). From day 35 to 42, early optic vesicles were cultured in DMEM/F12, 2% B27 supplement (50X), 1X NEAA, 1% Glutamax and 1% P/S. From day 42 to 70, ROs were moved to 24-well Cell Repellent Surface plates (Greiner Bio-One, Austria, cat#662970) cultured in retinal differentiation media 1 containing DMEM/F12, 2% B27 supplement (50X), 1X NEAA, 1% Glutamax, 10% Foetal Bovine Serum (ThermoFisher Scientific, USA, cat#A5256701), 100µM taurine (Sigma Aldrich, USA, cat#T8691) and 1% P/S. From day 70 to day 84, ROs were cultured in retinal differentiation media 2 containing DMEM/F12, 2% B27 supplement (50X), 1X NEAA, 1% Glutamax, 10% Foetal Bovine Serum, 100µM taurine, 1µM retinoic acid and 1% P/S. From day 84 to day 96, ROs were cultured in retinal differentiation media 3 containing DMEM/F12, 2% B27 supplement (50X), 1X NEAA, 1% Glutamax, 10% Foetal Bovine Serum, 100µM taurine, 0.5µM retinoic acid, 1X N2 supplement and 1% P/S. From day 96 onwards, ROs were cultured in retinal differentiation media 4 containing DMEM/F12, 2% B27 supplement (50X), 1X NEAA, 1% Glutamax, 10% Foetal Bovine Serum, 100µM taurine, 0.5µM retinoic acid, 1X N2 supplement, 1% Medium MW Hyaluronan (R&D Systems, USA, cat#GLR004) and 1% P/S.

### Retinal Pigment Epithelium Differentiation

Induced pluripotent stem cells were differentiated to RPE based on a protocol adapted from previous studies and initially characterized based on pigmentation and honeycomb cobblestone-like morphology. Briefly, iPSCs were cultured to 60-80% confluency and transitioned to RPE media 1 from day 0 to 7 containing DMEM high glucose, 20% Knockout Serum Replacement (KOSR; ThermoFisher Scientific, USA, cat#10828028), 1% P/S and 10mM Nicotinamide. From day 7-14, cells were cultured in RPE media 2 containing DMEM high glucose, 20% KOSR, 1% P/S and 100ng/mL Human Activin A (Miltenyi Biotec, Germany, cat#130-115-013). From day 14-42, cells were cultured in RPE media 3 containing DMEM high glucose, 20% KOSR, 1% P/S and 3µM CHIR99021 (Miltenyi Biotec, Germany, cat#130-103-926). Differentiating RPE was split at day 42 at a 1:5 ratio and cultured in RPE media 4 comprised of DMEM high glucose, 4% KOSR and 1% P/S. Cells were maintained an additional three weeks in RPE media 4 and split again at a 1:5 ratio. RPE was maintained in RPE media 4 until downstream analysis.

### RNA extraction and cDNA synthesis

Cells were lysed using a 1.5mL tube (JetBiofil, China, cat#CSP003002) in 1mL of TRI reagent (Sigma-Aldrich, USA, cat#T9424). RNA was extracted using the Direct-zol™ RNA MicroPrep (Zymo Research, USA, cat#ZR-R2060) or MiniPrep (Zymo Research, USA, cat#ZR-R2050) kits and treated with DNase I, Amplification Grade (ThermoFisher Scientific, USA, cat#18068015). RNA concentration was standardized and cDNA was synthesized using the High-Capacity cDNA Reverse Transcription Kit (ThermoFisher Scientific, USA, cat#4374966).

### Real time PCR and quantitative real time PCR analysis

Synthesized cDNA was used for RT-PCR and qRT-PCR analysis. For RT-PCR analysis, cDNA was amplified using PCRBio HS Taq Mix Red (PCR Biosystems Ltd, United Kingdom, cat#PB10.23-02) with appropriate primers according to manufacturer’s recommendations. Gene transcript levels were evaluated by qRT-PCR analysis using PowerTrack™ SYBR Green Master Mix as per the manufacturer’s recommendation (ThermoFisher Scientific, USA, cat#AB-A46109). Experiments were performed on the CFX96 Touch Real-Time PCR Detection System (BioRad Inc, USA). Genes with C t values & gt;32 were excluded from relative quantification analyses due to reduced assay sensitivity and increased variability. Each gene was amplified in triplicate, and the expression level of each gene was normalized to human GAPDH and ACTB as endogenous controls using the standard 2(ΔΔCT) method. All primers are listed in **Supplementary table 5.**

### Statistical analysis

Statistical analysis was performed by one-way analysis of variance (ANOVA) followed by two-sided Dunnett’s multiple comparisons test. Graphs were plotted using prism GraphPad.

## Results

### Arg>Ter PTCs: A major PTC subclass in IRDs and key target for ACE-tRNA therapy

Building upon our previously published IRD-PTC spectrum analysis ^28^, we have now extended the genetic dataset to include over 37,500 published IRD cases and used it to characterize the spectrum of PTCs more precisely. The analysis revealed that PTCs account for 18.3% of all IRD variants **(Figure 1A)**, establishing their substantial contribution to the mutational burden of IRDs. We identified 1649 unique published PTCs in 163 IRD genes. The distribution of these variants across genes **(Figure 1B)** revealed *USH2A* as the most frequently affected gene (11.5% of all PTCs), followed by *ABCA4* (8.8%), *RPGR* (5.8%), and *EYS* (5.4%). Analysis of nonsense codon origins **(Figure 1C)** showed that sense codons for Gln and Arg were the most common sources, each constituting 20.3% of all nonsense variants (335 unique PTCs for each type). This was followed by Glu (14.1%), Trp (13.3%), and Tyr (11.0%). Consequently, the most frequent nonsense codon was UAG (37.1%), followed closely by UGA (36.7%) and UAA (25.7%) **(Figure 1D)**.

**Figure 1:**
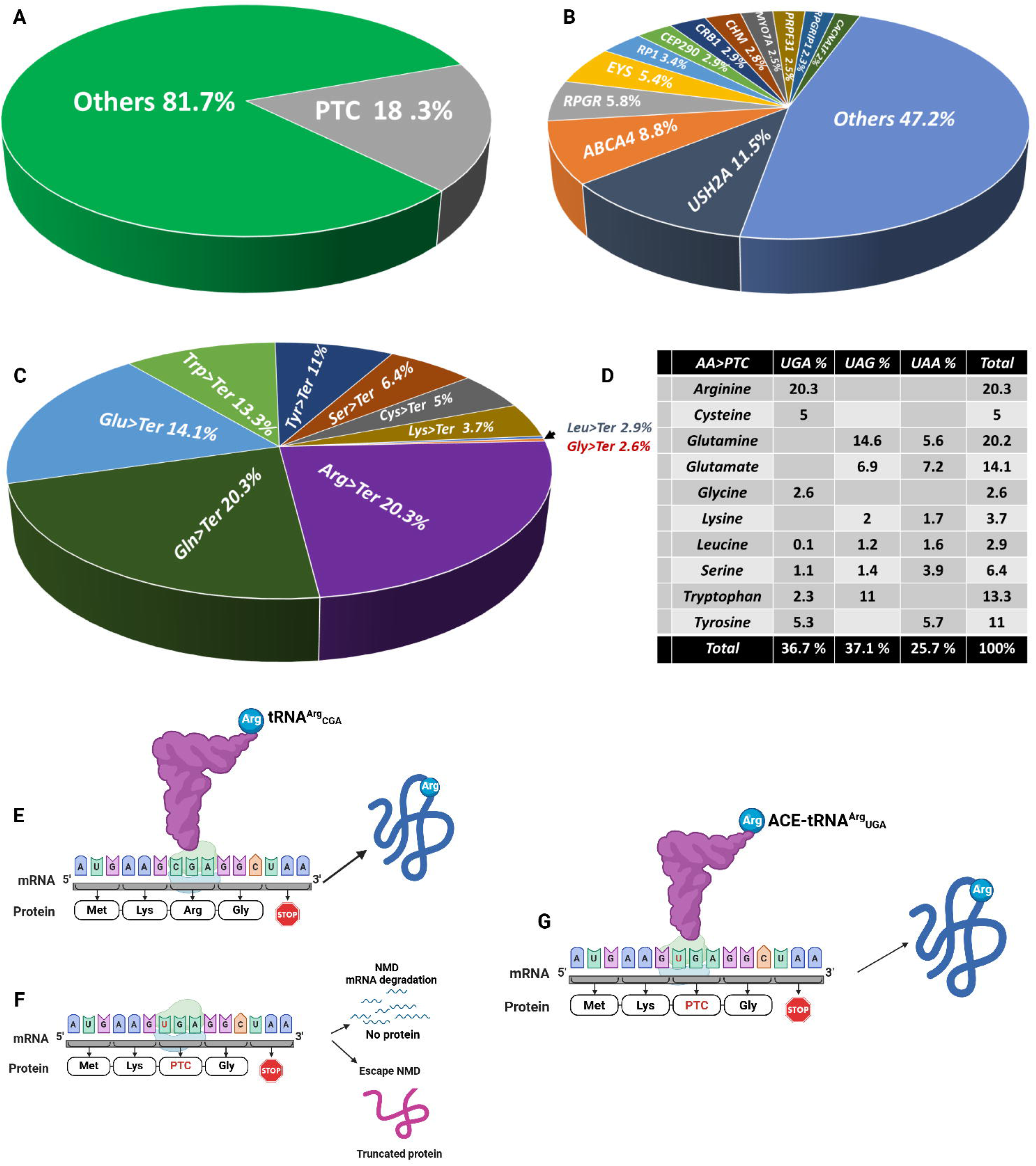
A survey of IRD-causing nonsense variants: **(A)** Proportion of PTC variants among all IRDs causing variants; **(B)** Frequency of PTC variants in different IRD-causing genes; **(C)** Percentage of different amino acids mutated to PTCs; **(D)** A detailed analysis of different sense codons mutated to different nonsense codons. **ACE-tRNA-mediated translational readthrough of premature termination codons: (E)** codon-anticodon base paring between mRNA and tRNA during translation; **(F)** Consequences of a PTC on mRNA; **(G)** Translational readthrough of a UGA PTC using ACE-tRNA^Arg^_UGA_

Further analysis revealed that although Arg>Ter PTCs represent only 20.3% of all nonsense variants, they affect 35% of IRD patients with PTCs. Consistently, 11 of the top 20 most frequent individual PTCs in this dataset are Arg>Ter, further confirming that this class is the most common and recurrent PTC type in IRD patients **((Supplementary Table 1).** Together, these findings establish Arg>Ter PTCs as a primary target for therapeutic intervention due to the potential translational impact. We therefore summarized that an ACE-tRNA therapy engineered for Arg>Ter PTCs is a compelling strategy, enabling a single ACE-tRNA to address the underlying cause in a significant portion of the IRD patient population **(Figure 1E-G)**.

### IRD genes display unique, gene-specific PTC signatures

We next analysed the PTC profiles of the four IRD genes with the highest number of PTCs **(Figure 2).** In *USH2A*, we identified 191 unique PTCs, with Trp, Gln, and Tyr codons being the most frequently mutated to PTCs. Among the three stop codons, UAG and UAA were the most common, followed by UGA **(Figure 2A).** In *ABCA4*, 146 unique nonsense variants were identified, where Trp, Gln, and Glu codons were most frequently mutated to PTCs. However, compared to *USH2A*, the nonsense codon distribution differed significantly, with UAG being the most abundant, followed by UAA and UGA **(Figure 2B)**. In *RPGR*, we identified 97 unique PTCs. Notably, over 50% of which arose from mutated Glu codons. The frequency of PTCs followed a similar trend to *USH2A* and *ABCA4*, with UAG and UAA being more common than UGA **(Figure 2C)**. In *EYS*, 90 unique nonsense variants were found. Interestingly, Cys codons were the most commonly mutated to PTCs, and UAA was the most frequent nonsense codon **(Figure 2D)**. Taken together, these observations highlight distinct, gene-specific nonsense variant signatures across different IRD genes.

**Figure 2:**
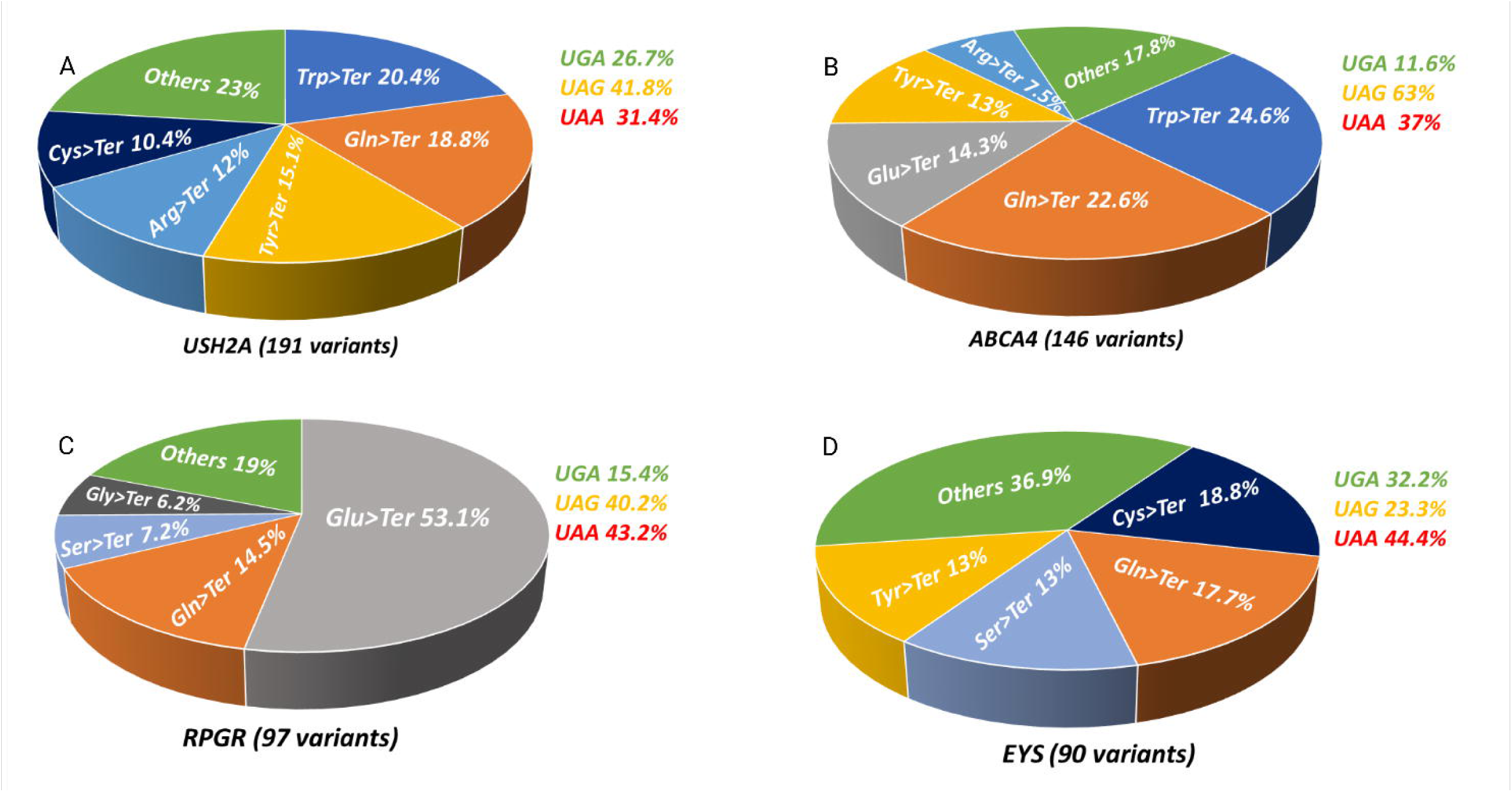
Gene specific nonsense variant signatures in IRD genes: **(A)** *USH2A*; **(B)** *ABCA4*; **(C)** *RPGR*; **(D)** *EYS*.

### Optimization of ACE-tRNA^Arg^ for efficient suppression of Arg>Ter nonsense variants

To improve the readthrough efficiency of Arg>Ter (UGA) nonsense codons, we aimed to design an optimized ACE-tRNA^Arg^ . We evaluated three distinct constructs: two previously characterized variants (V1 and V2) and a new construct, V3, which results from the combination of V2 and additional sequence elements **(Figure 3A & Supplementary table 3).** V1 contains an intron-less ACE-tRNA^Arg^ _UGA_ coding sequence under the control of a hU6 promoter (Albers et al., 2023) and V2 is driven by the same hU6 promoter but incorporates three specific nucleotide substitutions within the tRNA coding sequence (Albers et al., 2023). The V3 construct builds upon the modified tRNA sequence of V2 by integrating the recently reported 5’ upstream control element (UCE) and 3’ tiler sequence (TS), which have been shown to improve ACE-tRNA expression and suppression efficiency ^30^.

**Figure 3:**
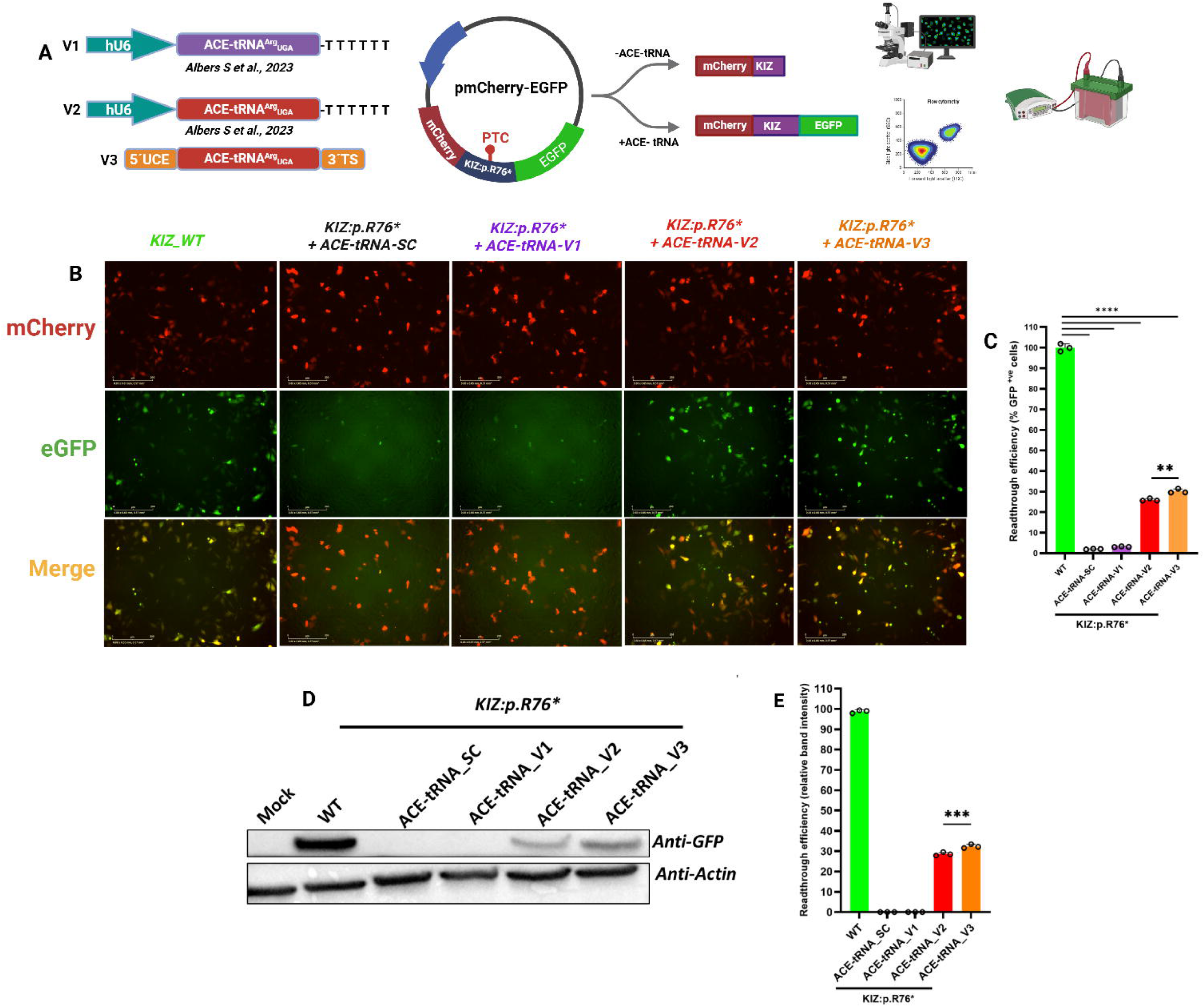
Optimization of ACE-tRNA^Arg^_UGA_ variants: **(A)** Schematic diagram of the experimental approach used to optimize the ACE-tRNA^Arg^; **(B)** Representative fluorescence microscopy images showing successful readthrough of KIZ:p.R73* PTC; **(C)** FACS analysis of ACE-tRNA_V3 readthrough efficiency (**=p<0.01); **(D)** Representative western blot image of GFP expression restoration following treatment with different ACE-tRNA^Arg^ variants; **(E)** Quantification of protein expression by Western blot (***=p<0.001). Data are mean presented as ± s.d from three biological replicates. Statistical analysis was performed by one-way analysis of variance (ANOVA) followed by two-sided Dunnett’s multiple comparisons test.

To test the efficiency of these three different ACE-tRNA^Arg^ molecules, we employed the *KIZ*:p.R76* PTC variant as a model system. This variant is one of the most common founder PTCs in the Jewish population causing autosomal recessive retinitis pigmentosa (arRP) ^31^. To this end, we cloned a 300-bp *KIZ* cDNA fragment encompassing 150 bp on either side of the PTC into the dual-reporter plasmid pmCherry-GFP. In this system, GFP expression occurs only upon successful readthrough of the PTC **(Figure 3A).** HeLa cells were co-transfected with each ACE-tRNA plasmid and the dual-reporter construct at a 1:2 mass ratio (0.5 µg of reporter plasmid and 1 µg of ACE-tRNA plasmid). Readthrough efficiency was then measured 48 hours post-transfection using a combination of fluorescent microscopy **(Figure 3B)**, flow cytometry, and western blot analysis. Flow cytometry revealed low V1 GFP expression (2.1% ± 3) and higher V2 (28.4 ± 2%) and V3 (33.2 ± 3%) GFP expression **(Figure 3C and S1)**, which were corroborated by western blot **(Figure 3D-E).** Given its performance, the V3 construct was used for all subsequent experiments.

### ACE-tRNA_V3 dosing yields high readthrough without compromising cell viability

For our preliminary screening, we employed a high dose (1 µg) of each ACE-tRNA variant to obtain a robust GFP signal. Under these conditions, ACE-tRNA-V2 and ACE-tRNA-V3 caused a notable decrease in cell confluence (Data not shown), implying that higher doses of these variants may compromise cellular health. To define the therapeutic window for V3, we performed a concentration-response readthrough assay by titrating the V3 plasmid (50-500ng) against a constant 500ng of the *KIZ* reporter plasmid, with the total transfected DNA equalized using empty plasmid filler. The results revealed 200ng of V3 as the optimal dose, producing a maximal readthrough efficiency of 39 ±2% **(Figure 4A).** A gradual decline in efficiency was observed at higher concentrations (400-500ng). This toxicity-associated impairment is further highlighted by a direct comparison: while the 200ng dose yielded 39% efficiency, a dose of 1000 ng in our initial comparison experiment yielded only 33% **(Figure 3B-E).** This significant reduction in performance at a five-fold higher dose underscores that V3’s therapeutic benefit is directly counteracted by its toxicity beyond an optimal concentration.

**Figure 4:**
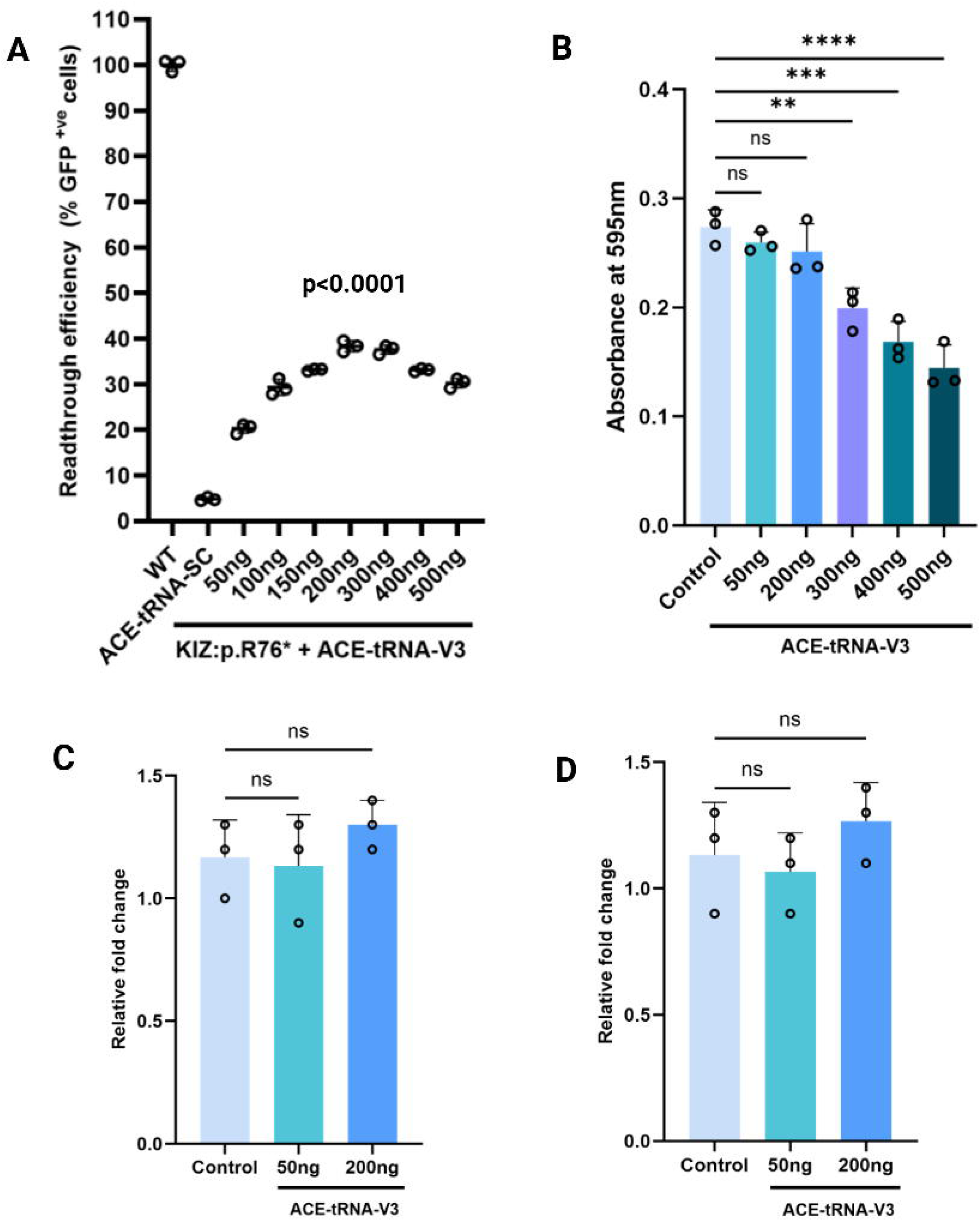
ACE-tRNA^Arg^_UGA_-V3 shows effective readthrough with minimal toxicity: **(A)** Readthrough efficiency of the KIZ:p.R76* reporter as a function of ACE-tRNA^Arg^-V3 plasmid dosage (50-500 ng) (P < 0.0001); **(B)** Cell viability assessed by MTT assay following transfection with increasing amounts of the ACE-tRNA^Arg^ -V3 construct (0-500 ng). **(C)** qPCR quantification of BIP and CHOP cDNA from HeLa cells transfected with ACE-tRNA^Arg^_UGA_-V3. Data are presented as mean ± s.d. from three biological replicates. Statistical analysis was performed by one-way analysis of variance (ANOVA) followed by two-sided Dunnett’s multiple comparisons test (NS=non-significant; **=p<0.01; ***=p<0.001; ****=p<0.0001).

To corroborate these findings, we performed an MTT cell viability assay, which confirmed high toxicity at doses of 300 ng and above, while doses between 50-200 ng showed minimal effect on viability **(Figure 4B)**. Additionally, we measured the gene expression levels of C/EBP homologous protein (CHOP) and Binding immunoglobulin Protein (BiP), markers of endoplasmic reticulum stress and activation of the unfolded protein response induced by global readthrough and production of C-terminal extended proteins. At the optimal 200 ng dosage, we did not observe any significant increased expression of these genes **(Figure 4C-D)**.

### ACE-tRNA**-**V3 efficiently suppresses multiple prevalent IRD nonsense variants

To evaluate the broader therapeutic potential of ACE-tRNA-V3 using the established optimal dose of 200ng, we tested ACE-tRNA_V3 against 13 pathogenic Arg>Ter variants (Figure 5A-C). Readthrough efficiency ranged from 14.8% to 86.1%, stratifying into three tiers: high (>50%: FAM161A:p.R523*- 86.1%; USH2A:p.R737*- 70.7%; RP1:p.R677*- 67.6%; RP1L1:p.R658*- 68.2%; PRCD:p.R22*- 62.5%), medium (35–50%: ABCA4:p.R2149*-48.7%; FAM161A:p.R437*- 47.5%; CNGA1:p.R28*- 45.6%; KIZ:p.R76*- 39%), and low (<35%: CERKL:p.R283*- 31.6%; RP1:p.R1933*,- 30.1%; ABCA4:p.R2030*- 28.7%; ABCA4:p.R681*- 14.8%).

**Figure 5:**
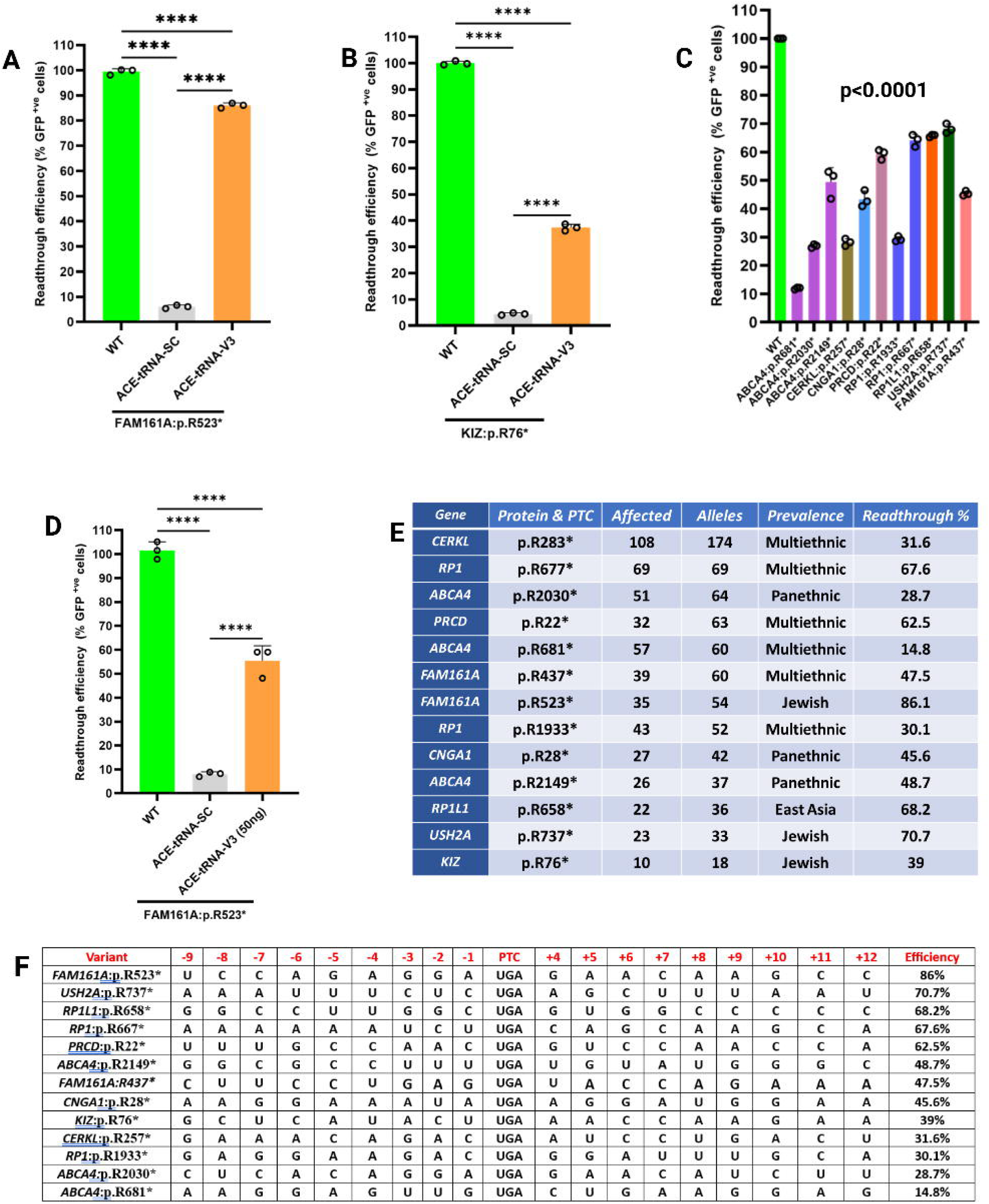
ACE-tRNA^Arg^_UGA_-V3 induces variable nonsense suppression across a panel of IRD-associated variants: **(A-B)** Flow cytometry analysis of readthrough efficiency for the founder variants KIZ:p.R76* (A) and FAM161A:p.R523* (B) following treatment with ACE-tRNA^Arg^ -V3 (200 ng) (****=p<0.0001). **(C)** Quantification of readthrough efficiency for UGA an extended panel of 11 additional pathogenic variants, measured by flow cytometry (****=p<0.0001). **(D)** FACS analysis showing readthrough efficiency of FAM161A:p.R523* with 50 ng of ACE-tRNA^Arg^_UGA_-V3 (reporter: 500 ng) (****=p<0.0001). Data are presented as mean ± s.d. from three biological replicates. Statistical analysis was performed by one-way analysis of variance (ANOVA) followed by two-sided Dunnett’s multiple comparisons test. **(E)** Corresponding table of all tested variants, listing gene, variant, reported affected individuals, allele count, population prevalence, and quantified readthrough percentage. **(F)** Table showing the +4 nucleotide and suppression efficiency (%) for 13 Arg>Ter variants.

To investigate the basis for this variability, we predicted RNA secondary structure upstream of the PTC. Low-readthrough ABCA4:p.E683* formed a highly stable structure (MFE = – 25.50 kcal/mol), whereas high-readthrough FAM161A:p.R523* and USH2A:p.R737* exhibited less stable structures (MFE = –18.60 and –13.10 kcal/mol, respectively) (Figure S2). This suggests that local mRNA stability may impede ACE-tRNA accessibility.

The exceptionally high readthrough of FAM161A:p.R523* prompted us to test whether lower doses could maintain efficacy. Indeed, 50 ng ACE-tRNA_V3 achieved 60% readthrough (compared to 86% at 200 ng), demonstrating that highly permissive contexts require lower drug concentrations (Figure 5D). This demonstrates that the optimal therapeutic dose of ACE tRNA may be variant dependent and can be reduced for highly amenable sequence contexts. A summary of the number of cases and readthrough efficacy for each variant is shown in Figure 5E. We next examined whether the +4-nucleotide downstream of the PTC influences ACE-tRNA readthrough efficiency, as this position has been reported to affect PTC124-dependent readthrough. By comparing the +4-sequence context across the 13 tested variants, we observed no clear trend of altered suppression levels **(Figure 5F)** suggesting that the +4-position alone does not predict ACE-tRNA readthrough efficiency.

### ACE-tRNA^Arg^_UGA__V3 restores wild-type PRCD expression and intracellular trafficking

Biallelic pathogenic variants in *PRCD* are known to cause arRP. The p.R22* nonsense variant, which is highly prevalent in an isolated Arab-Muslim village in Northern Israel ^32^, is a favourable candidate for PTC suppression due to the small open reading frame of *PRCD* and well-defined subcellular localization of PRCD to the endoplasmic reticulum and Golgi ^33^. To leverage these features, we cloned the entire *PRCD* cDNA into a C-terminal eGFP reporter plasmid. This approach differs from the dual mCherry-GFP reporter system used for other variants, as it omits the N-terminal mCherry tag to avoid any potential interference with the native localization signal **(Figure 6A).**

**Figure 6:**
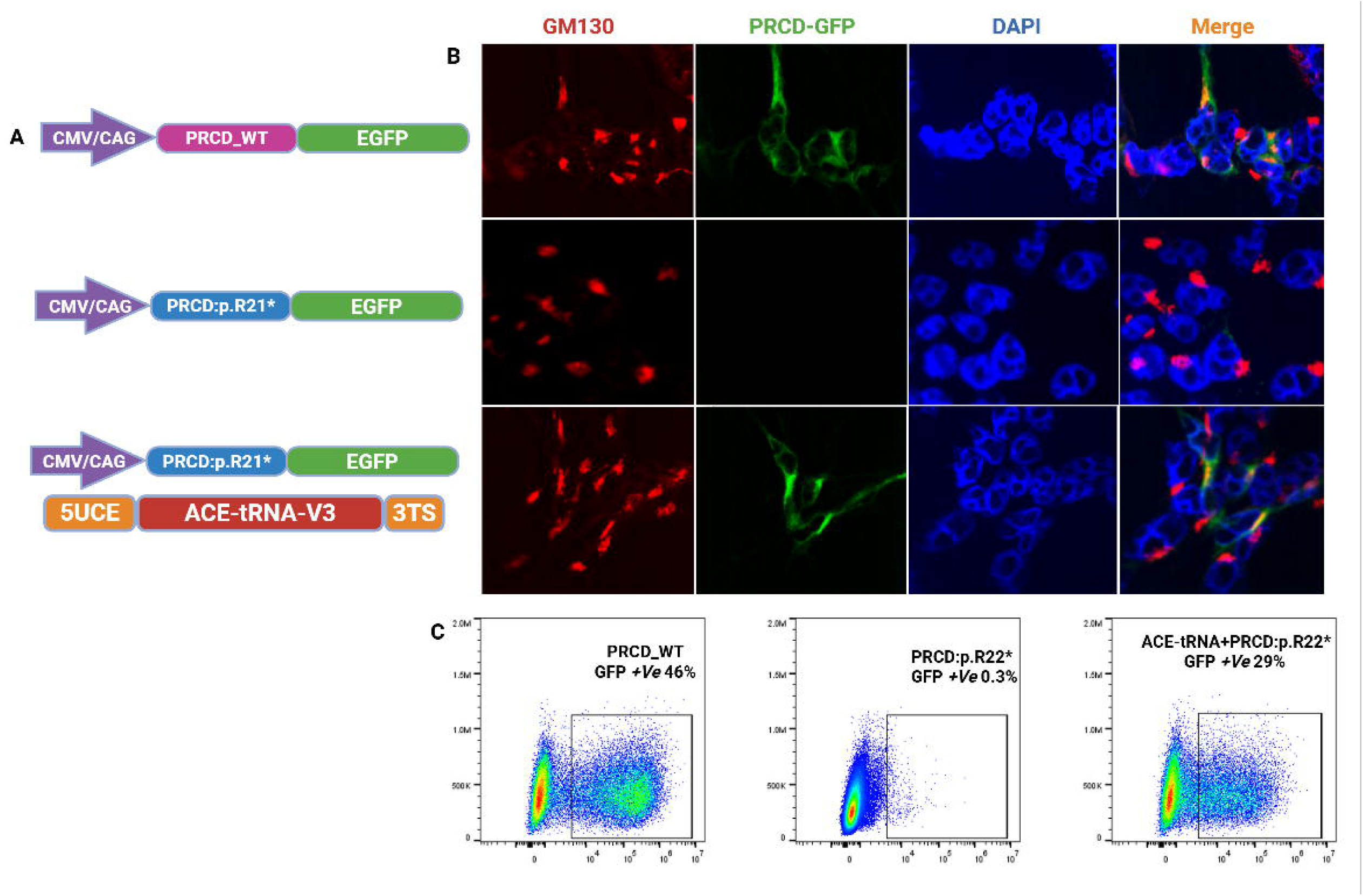
ACE-tRNA_V3 rescues PRCD:p.R21* expression and localization: **(A)** Schematic representation of the plasmid constructs used in this assay; **(B)** Representative immunofluorescence images of HeLa cells transfected with PRCD-WT-eGFP (top), PRCD:p.R21*-eGFP (middle), or PRCD:p.R21*-eGFP + ACE-tRNA_V3 (bottom). PRCD-EGFP (green), GM130 (red), DAPI (blue). Scale bar = 10 µm; **(C)** Representative FACS histograms showing percentage of eGFP positive HeLa cells transfected with the indicated constructs. The percentage of eGFP-positive cells is indicated for each condition.

We then co-transfected HeLa cells with either wild-type or p.R22* mutant constructs alongside ACE-tRNA^Arg^ _V3 and performed high-resolution fluorescent microscopy using an antibody against the Golgi marker GM130. Immunofluorescence analysis revealed that ACE-tRNA^Arg^ _V3 treatment efficiently restored the subcellular localization of the mutant PRCD to the Golgi, demonstrating robust co-localization with Golgi marker GM130. This result confirms that ACE-tRNA therapy not only facilitates readthrough of the PTC but also rescues the functional trafficking of PRCD **(Figure 6B)**, while FACS quantification of the treated cell population demonstrated a PTC readthrough efficiency of 62% **(Figure 6C).**

### ACE-tRNA_V3 mediates nonsense readthrough in 661W photoreceptor-like cells

Having established the efficacy of ACE-tRNA_V3 in HeLa cells, we next sought to validate its functional activity in a photoreceptor-relevant cellular context. To this end, we employed 661W cells, an immortalized mouse cone-like photoreceptor cell line that show some key features of native retinal neurons ^34,35^. Given the difficulty of transfecting 661W cells compared to HeLa cells, we generated a single plasmid construct, pDS-003, which contains both the ACE-tRNA_V3 expression cassette and an mCherry-FAM161A reporter of either wild-type or carrying the p.R523* nonsense variant. This approach eliminates the need for dual transfection and ensures consistent delivery of both components.

661W cells were transfected with pDS-003 constructs encoding either mCherry-FAM161A-WT or mCherry-FAM161A:p.R523*. Readthrough efficiency was assessed 48 hours post-transfection via Incucyte live-cell imaging. Cells expressing the WT reporter displayed robust mCherry fluorescence, confirming proper expression. In contrast, cells expressing the mutant reporter alone showed no detectable mCherry signal, whereas those co-expressing the mutant reporter with ACE-tRNA_V3 exhibited strong fluorescence **(Figure 7A).** These results demonstrate efficient readthrough of the FAM161A:p.R523* nonsense variant in this photoreceptor-like cell line. Importantly, these findings demonstrate that ACE-tRNA_V3 retains its nonsense suppression activity in a cellular environment closely resembling native retinal neurons, underscoring its translational potential for IRD therapy.

**Figure 7:**
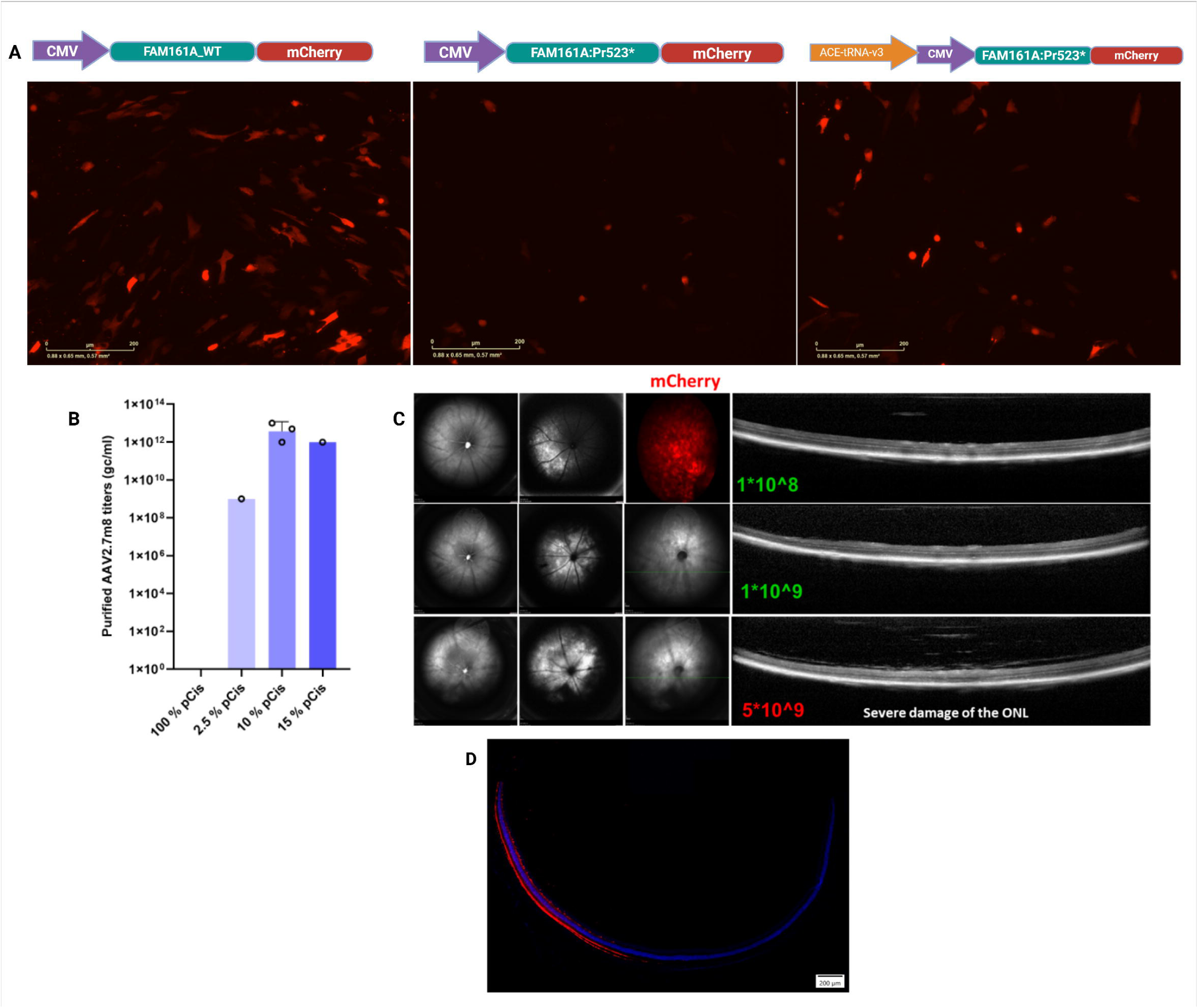
ACE-tRNA_V3 mediates nonsense readthrough in 661W cells and AAV2/7m8 packaging enables safe retinal delivery: **(A)** Fluorescence microscopy images of 661W cells transfected with pDS-003 containing mCherry-FAM161A WT (left), mCherry-FAM161A:p.R523* mutant alone (middle), or FAM161A:p.R523* mutant with ACE-tRNA_V3 (right). **(B)** Bar graph showing AAV2/7m8 titers obtained using different cis-plasmid percentages (2.5%, 10%, and 15%) during low-cis triple transfection. **(C)** Representative OCT images of wild-type mouse retinas 5 weeks post-subretinal injection of AAV2/7m8-ACE-tRNA_V3 at 1×10, 1×10, and 5×10 GC/eye. **(D)** Immunohistochemistry of retinal sections showing mCherry expression (red) on the injected side, confirming viral transduction at 1×10 GC/eye. Nuclei counterstained with DAPI (blue).

### Low-cis triple transfection enables AAV2/7m8 packaging of ACE-tRNA_V3 for safe in vivo retinal delivery and efficient RPE and ROs transduction in vitro

Packaging of ACE-tRNA constructs into AAV particles is inherently challenging, as using 100% cis plasmid results in poor or undetectable viral titers. Low-cis triple transfection has emerged as an effective alternative to overcome this bottleneck ^36^. However, the optimal cis-plasmid percentage varies across AAV serotypes and has not been established for AAV2/7m8 ^36^, a capsid variant with superior photoreceptor tropism ^26,37^. To address this, we tested three different cis-plasmid percentages (2.5%, 10%, and 15%) during AAV2/7m8 production using the triple transfection method. Among the conditions tested, 10% cis plasmid yielded the optimal packaging efficiency, producing viral titers consistently ranging from 1×10¹² to 1×10¹³ genome copies per mL (GC/mL). In contrast, both 2.5% and 15% cis conditions resulted in substantially lower titers **(Figure 7B).**

To confirm that the packaged AAV2/7m8-ACE-tRNA_V3 retains functional activity, we tested the virus in HeLa cells transfected with the mCherry-KIZ:p.R76*-GFP reporter. HeLa cells were co-transfected with the reporter plasmid and subsequently transduced with AAV2/7m8-ACE-tRNA_V3 at a multiplicity of infection corresponding to the previously established safe titer. Readthrough efficiency was assessed by fluorescence microscopy. In cells transduced with AAV2/7m8-ACE-tRNA_V3, robust GFP expression was observed, indicating successful readthrough of the KIZ:p.R76* nonsense variant. In contrast, control cells (reporter only, without virus) exhibited no detectable GFP fluorescence **(Supplementary figure S3).** These findings confirm that AAV2/7m8-ACE-tRNA_V3 retains nonsense suppression activity following packaging and transduction, validating its functionality for subsequent in vivo studies.

Having established a robust AAV2/7m8 production method and confirmed viral functionality, we next evaluated the safety of AAV2/7m8-ACE-tRNA_V3 *in vivo*. Wild-type mice received subretinal injections of the virus at three different titers: 1×10, 1×10, and 5×10^9^ GC/eye. Retinal toxicity was assessed in vivo by optical coherence tomography (OCT) at 5 weeks post-injection (N=5, 5 eyes for each titer). OCT analysis revealed that the 1×10 titer was well-tolerated, with preserved retinal morphology **(Figure 7C).** In contrast, the higher 5×10 titer induced retinal toxicity, characterized by thinning of the outer nuclear layer (photoreceptors) and disruption of photoreceptor outer segments. Based on these findings, 1×10 GC/eye was established as the maximum safe dose **(Figure 7C-D).**

Finally, to assess transduction efficiency in a human-relevant cell type, we transduced WT iPSC-derived RPE cells and day 180 ROs with AAV2/7m8-ACE-tRNA_V3 at the optimal titer. AAV2/7m8-ACE-tRNA_V3 transduced both RPE cells and ROs efficiently, demonstrating that AAV2/7m8 delivers ACE-tRNA_V3 to both photoreceptors and the RPE cells **(Figure S4).** Together, these findings establish a viable production and pre-clinical delivery platform for AAV2/7m8-ACE-tRNA_V3, supporting its continued development for IRD therapy.

## Discussion

While recent studies have provided important proof-of-concept for ACE-tRNA therapy in the mouse retina^24,25^, their experimental designs were necessarily focused on establishing mechanistic feasibility rather than targeting clinically relevant nonsense variants. In one case ^24^, a murine variant (RPE65:p.Arg44Ter) that is not relevant for humans was studied while the other ^25^ examined an ultra-rare nonsense variant (KCJ13:p.Trp53Ter) with very limited patient prevalence. Therefore, a critical next step toward clinical translation is the systematic identification of nonsense variant subclasses that affect substantial patient populations and the development of therapies tailored to them. Here, we address this need by analyzing over 37,500 IRD patients to define the targetable nonsense variant spectrum. The finding that Arg>Ter variants constitute 20.3% of all IRD nonsense variants yet affect 35% of nonsense-variant patients highlights their disproportionate clinical burden. This discrepancy arises because many Arg>Ter variants recur across multiple unrelated patients, as evidenced by 11 of the top 20 most frequent individual nonsense variants being Arg>Ter. These data establish Arg>Ter as the priority target for nonsense suppression therapy in IRDs.

Our genetic analysis of over 37,500 IRD patients across 163 genes and 1,650 unique nonsense variants provides a valuable data-driven framework for the field. This comprehensive compendium serves as a reference for prioritizing therapeutic targets, informing clinical trial recruitment, and advancing mutation class-based nonsense suppression approaches beyond our ACE-tRNA strategy.

Beyond the global predominance of Arg>Ter variants, our analysis revealed that individual IRD genes display unique, gene-specific nonsense variant signatures. In *RPGR*, over 50% of nonsense variants arise from Glu codons (due to the unique amino acid sequence of ORF15), suggesting that an ACE-tRNA targeting Glu>Ter would address most nonsense variants in this gene. In *EYS*, Cys codons are the most frequently mutated sense codons, while UAA is the most common PTC. Given that UAA is the least readthrough-responsive stop codon (Albers et al., 2023b; Ko et al., 2022b)(Albers et al., 2023b; Ko et al., 2022b), alternative strategies may be required for *EYS*, such as using multiple copies of the ACE-tRNA (e.g., 4x or 8x expression cassettes) or engineering highly potent ACE-tRNA variants with enhanced activity ^39^. Collectively, these gene-specific signatures provide a practical roadmap for prioritizing development of ACE-tRNAs to meet clinical needs.

Having established the global and gene-specific nonsense variant landscape, we next developed and optimized an ACE-tRNA targeting the most prevalent subclass Arg>Ter UGA codons. Our optimization studies reveal several insights for ACE-tRNA design. First, the modest improvement of V3 over V2, compared to the large difference between V2 and V1, suggests that the tRNA body sequence is the primary determinant of readthrough efficiency. This finding aligns with recent work showing that anticodon loop and D-arm modifications significantly improve ACE-tRNA efficiency (Akyuz et al., 2025; Albers et al., 2023c; Ren et al., 2025b). Second, the narrow range of non-toxic ACE-tRNA plasmid dosage suggests that the same properties that enable efficient readthrough may also increase off-target suppression of natural stop codons or induce cellular stress. Notably, the observation that FAM161A:p.R523* achieved near-maximal readthrough at only 50ng demonstrates that highly permissive sequence contexts can circumvent this toxicity. From a translational perspective, these findings underscore the importance of careful dose optimization for each target variant where patient genotypes may inform ACE-tRNA dosage.

The variability in readthrough efficiency across different Arg>Ter variants ranging from 15% to 86% was striking, given that all target the same UGA PTC. This observation reinforces the notion that PTC suppression is governed by more than just the stop codon identity. Local sequence context, including mRNA secondary structure and translational kinetics, profoundly influences how efficiently a near-cognate tRNA can outcompete release factors at the ribosome ^24,41^. Interestingly, the +4-nucleotide position, a well-established determinant of PTC124 activity ^42^, did not predict ACE-tRNA efficiency in our panel. This indicates that the sequence determinants of ACE-tRNA readthrough differ from those reported for small-molecule readthrough agents. Supporting this, our secondary structure analysis showed that low-readthrough variants formed more stable RNA structures upstream of the PTC, suggesting that local mRNA structure and consequently translational velocity may be key determinants of ACE-tRNA readthrough efficiency. From a clinical development perspective, the composition of our variant panel is noteworthy: nine of the 13 tested alleles are pan-ethnic, reported in IRD patients worldwide, while the remainder are founder mutations in the Jewish population. This diversity suggests that an ACE-tRNA therapy targeting Arg>Ter UGA PTCs would have global applicability, addressing both common recurrent nonsense variants and population-specific alleles. Beyond quantitative readthrough, we show that ACE-tRNA_V3 restores native PRCD localization to the Golgi. PRCD mislocalization (Murphy & Kolandaivelu, 2016; Spencer et al., 2016), is linked to RP, and its rescue confirms that the readthrough protein is properly trafficked. Thus, ACE-tRNA_V3 achieves functional restoration for PRCD:p.R22*. Future work in patient-derived ROs and mouse models will further validate these findings.

Efficient readthrough by ACE-tRNA_V3 in 661W photoreceptor-like cells confirms its activity in a retinal-like context, consistent with previous reports that ACE-tRNAs optimized in HEK293 cells perform similarly across tissue ^24,25,29,45^. These findings support the translational potential of ACE-tRNA_V3 for IRD therapy, though in vivo validation is warranted.

A major hurdle for translating ACE-tRNA therapy to the clinic is the difficulty of packaging these small, structured RNA molecules into AAV capsids. Using 100% cis plasmid resulted in poor viral yields, consistent with previous reports ^36^. To overcome this, we employed a low-cis triple transfection approach, testing three different cis-plasmid percentages for AAV2/7m8, a capsid variant with superior photoreceptor tropism. The finding that 10% cis plasmid yielded titers of 1×10¹²–1×10¹³ GC/mL establishes a viable production method for this serotype. A titer of 1×10 GC/eye did not cause retinal abnormalities in mice as determined by in vivo OCT imaging at 5 weeks post-injection. Furthermore, AAV2/7m8-ACE-tRNA_V3 efficiently transduced iPSC-derived RPE cells, demonstrating its utility for targeting both photoreceptors and the RPE. These packaging and safety data lay the groundwork for efficacy studies in IRD mouse models.

Several limitations of the current study warrant consideration. First, while we demonstrate activity in 661W cells, these are immortalized cells that may not fully recapitulate the complex retinal environment. Second, our in vivo data are limited to safety in wild-type mice; efficacy in disease models remains to be tested. Finally, the long-term safety and durability of AAV-mediated ACE-tRNA expression require further investigation.

In conclusion, this study provides a roadmap from large-scale patient genetics to preclinical nonsense suppression therapy for IRDs. We identified Arg>Ter nonsense variants as the dominant targetable subclass, affecting the largest share of IRD patients with nonsense mutations. Guided by this insight, we optimized ACE-tRNA (V3) that efficiently suppresses these PTCs, rescues functional protein localization, and can be delivered safely via AAV2/7m8. While efficacy in IRD preclinical models remains to be tested, this work establishes a precision medicine strategy that leverages genomic data to target the most common nonsense variant class with a single therapeutic molecule.

## Data availability statement

All data and experimental parameters related to this paper are available in the main text and Supplemental Figures. All raw data from this study are available from the corresponding author upon reasonable request.

## Acknowledgments

The authors would like to thank The ELSC Vector Core Facility (EVCF) at the Hebrew University of Jerusalem for their assistance with AAV packaging. The 661W photoreceptor cell line was generously provided by Dr Muayyad Al-Ubaidi (Department of Cell Biology, University of Oklahoma Health Sciences Center, Oklahoma City, OK, USA).

## Author contributions

A.S.S, J.E, A.S, H.K, MS, CM, AO, and SK performed experiments and interpreted the data, A.S.S, D.S, E.B conceived the experiments, A.S.S wrote the manuscript. D.S. supervised the study, provided funding, and edited the manuscript.

