## supplementary file for "Single anticodon-edited tRNA therapy targeting highly prevalent Arg>Ter premature termination codons causing inherited retinal diseases"

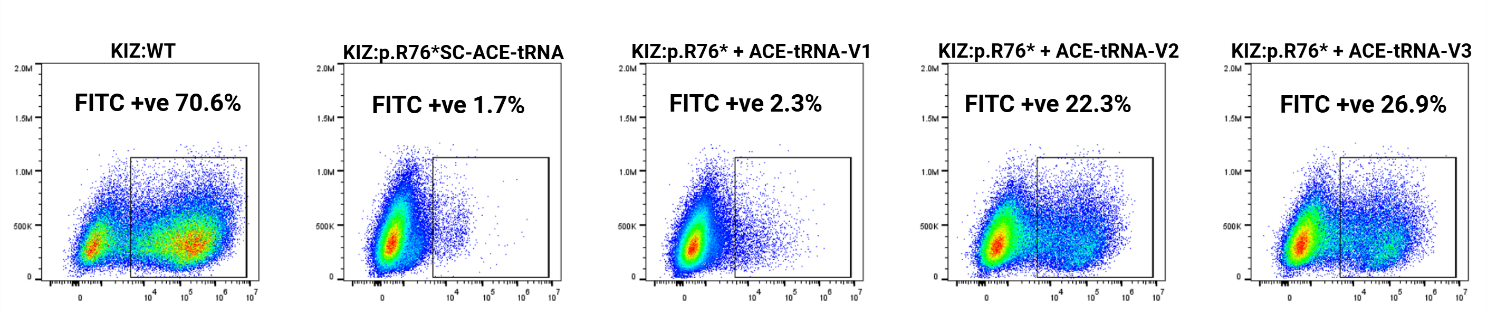

**Figure S1:** Representative FACS histograms showing GFP fluorescence intensity in HeLa cells transfected with the indicated constructs. The percentage of GFP-positive cells is indicated for each condition.

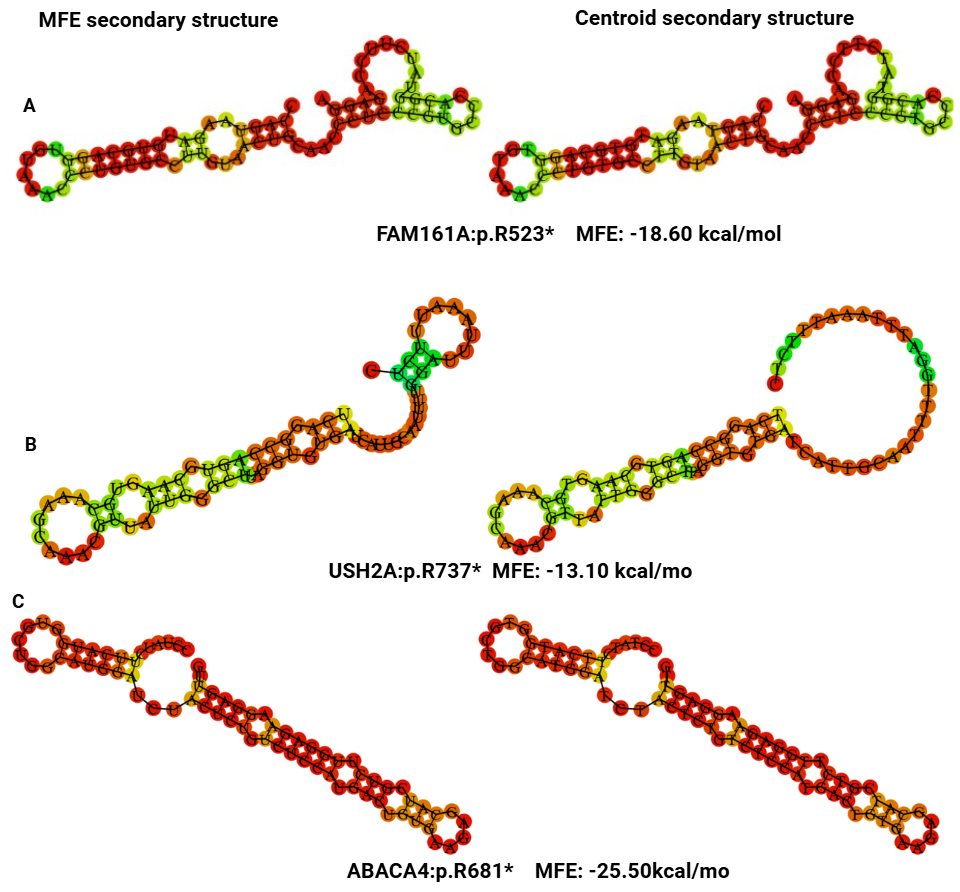

**Figure S2: Predicting the mRNA secondary structure using RNAfold web server** ([**http://rna.tbi.univie.ac.at/cgi-bin/RNAWebSuite/RNAfold.cgi**](http://rna.tbi.univie.ac.at/cgi-bin/RNAWebSuite/RNAfold.cgi)**): (A)** FAM161A:p.R523*; **(B)** USH2A:p.R737*; **(C)** ABCA4:p.R681*

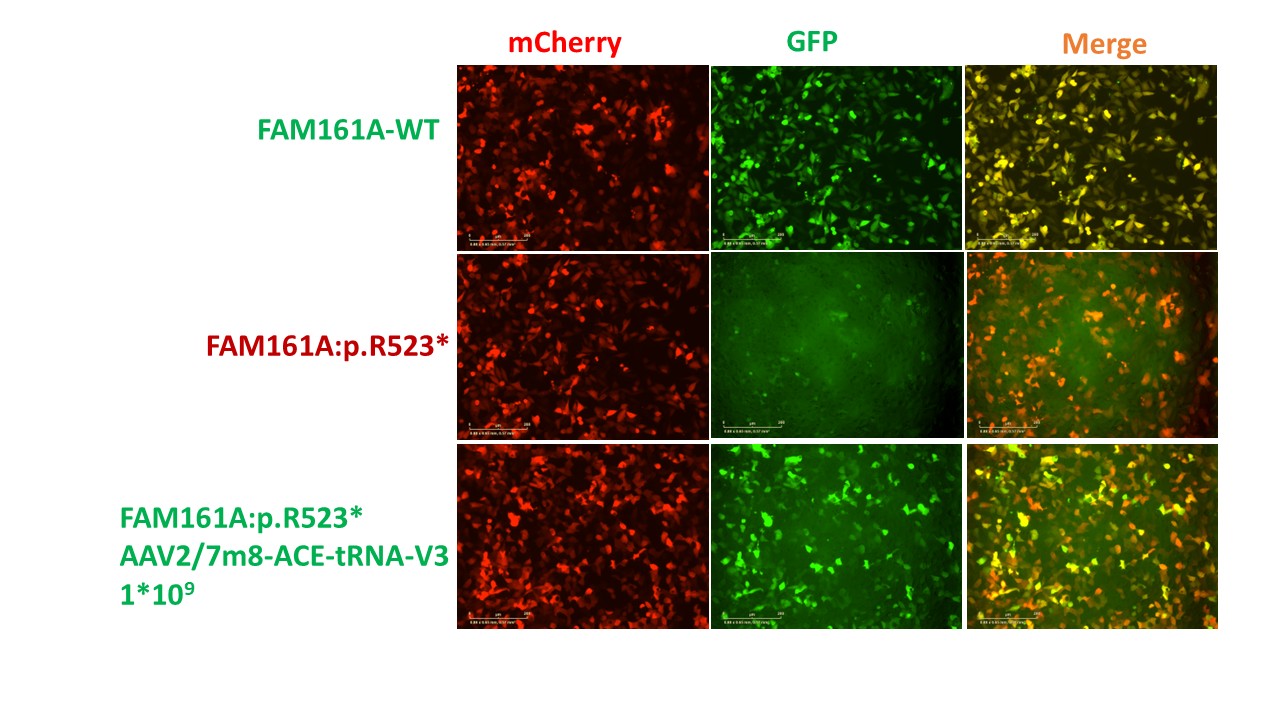

**Figure S3: FAM161A:p.R523* readthrough in HeLa cells transduced with AAV2/7m8-ACE-tRNA_V3;** Representative fluorescence images from Incucyte live cell imaging **(A):** FAM161A-WT; **(B)** FAM161A:p.R523*; **(C):** FAM161A:p.R523* with AAV2/7m8-ACE-tRNA_V3

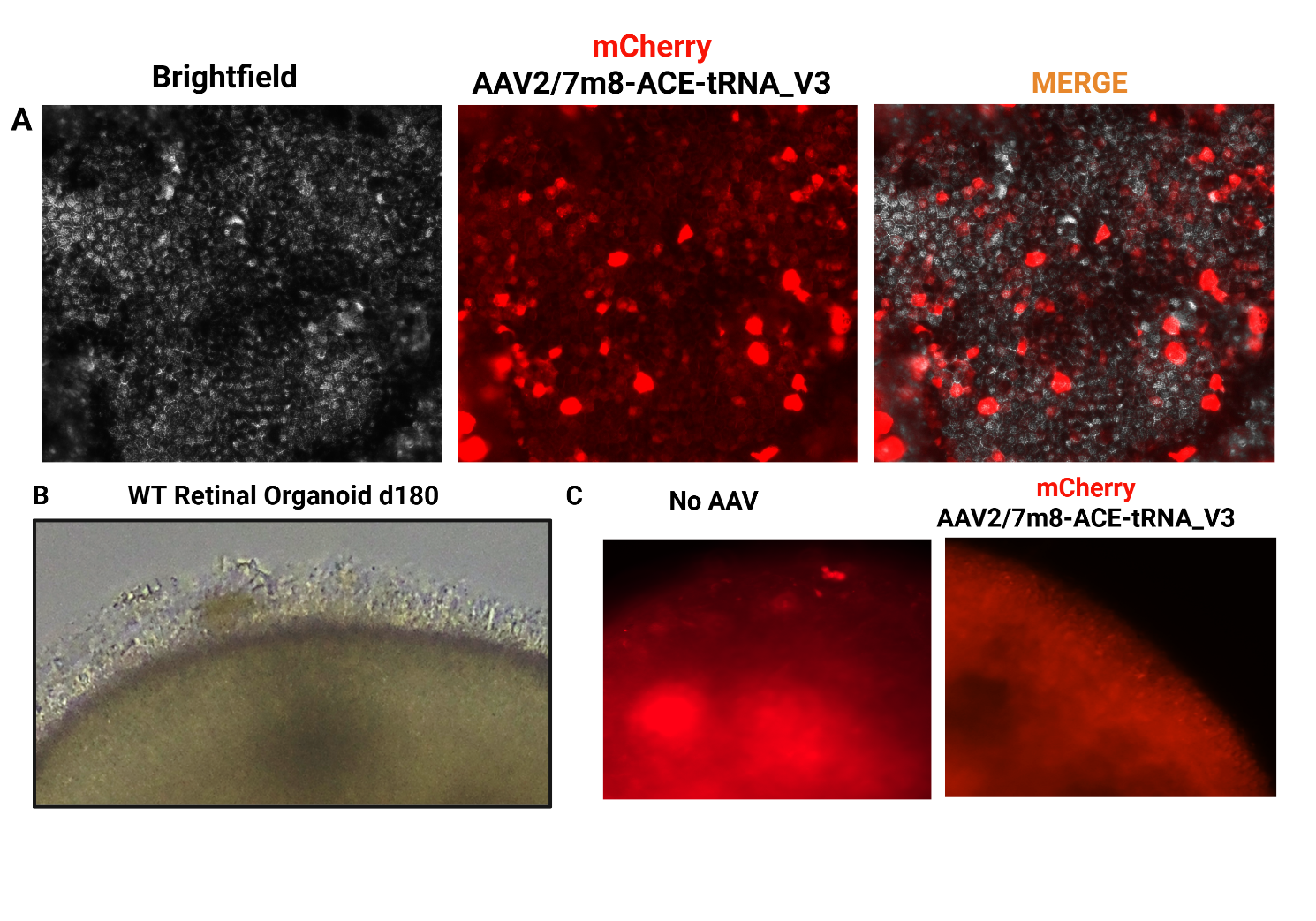

**Figure S4: AAV2/7m8 efficiently transduces both RPE and Retinal organoids: A.** Fluorescence microscopy images of WT iPSC-derived RPE cells transduced with AAV2/7m8-ACE-tRNA_V3 showing mCherry expression (red). **B.** Fluorescence microscopy images of WT iPSC-ROs transduced with AAV2/7m8-ACE-tRNA_V3 showing mCherry expression (red).

**Supplementary Table 1: Top 25 nonsense variants in IRD patients**

| **Gene** | **NM_ID** | **cDNA** | **Protein** | **PTC Type** | **Sum affected_total** | **Sum alleles_total** |
| --- | --- | --- | --- | --- | --- | --- |
| EYS | NM_001142800.2 | c.8805C>A | p.Y2935* | UAA | 188 | 228 |
| TULP1 | NM_003322.6 | c.901C>T | p.Q301* | UAG | 94 | 187 |
| CERKL | NM_001030311.3 | c.847C>T | p.R283* | UGA | 108 | 174 |
| USH2A | NM_206933.4 | c.11864G>A | p.W3955* | UAG | 101 | 112 |
| AIPL1 | NM_014336.5 | c.834G>A | p.W278* | UGA | 70 | 110 |
| ABCA4 | NM_000350.3 | c.4234C>T | p.Q1412* | UAG | 74 | 74 |
| RP1 | NM_006269.2 | c.2029C>T | p.R677* | UGA | 69 | 69 |
| ABCA4 | NM_000350.3 | c.6658C>T | p.Q2220* | UAG | 52 | 66 |
| ABCA4 | NM_000350.3 | c.6088C>T | p.R2030* | UGA | 51 | 64 |
| PRCD | NM_001077620.3 | c.64C>T | p.R22* | UGA | 32 | 63 |
| ABCA4 | NM_000350.3 | c.2041C>T | p.R681* | UGA | 57 | 60 |
| FAM161A | NM_001201543.2 | c.1309A>T | p.R437* | UGA | 39 | 60 |
| TRPM1 | NM_002420.6 | c.880A>T | p.K294* | UAG | 28 | 56 |
| FAM161A | NM_001201543.2 | c.1567C>T | p.R523* | UGA | 35 | 54 |
| RP1 | NM_006269.2 | c.5797C>T | p.R1933* | UGA | 43 | 52 |
| RP1 | NM_006269.2 | c.1625C>G | p.S542* | UAA | 29 | 47 |
| CNGA1 | NM_001379270.1 | c.82C>T | p.R28* | UGA | 27 | 42 |
| ABCA4 | NM_000350.3 | c.6445C>T | p.R2149* | UGA | 26 | 37 |
| RP1L1 | NM_178857.6 | c.1972C>T | p.R658* | UGA | 22 | 36 |
| EYS | NM_001142800.2 | c.7919G>A | p.W2640* | UAG | 31 | 36 |
| USH2A | NM_206933.4 | c.2209C>T | p.R737* | UGA | 23 | 33 |
| CRB1 | NM_201253.3 | c.2401A>T | p.K801* | UAG | 29 | 32 |
| RPGR | NM_001034853.2 | c.259G>T | p.E87* | UAA | 32 | 32 |
| PCDH15 | NM_033056.4 | c.733C>T | p.R245* | UGA | 17 | 31 |

**Supplementary Table 2: cDNA fragments cloned into mCherry-GFP reporter plasmids (Both wildtype and mutant sequence are cloned for feasibility only mut sequences are shown)**

| *FAM161A*:p.R523* | TTGGCAGACATCGAAGCAGATGAAGAAAATTTAAAAGAAACACGTTGGCCTTATTTGTCTCCAAGGCGTAAGTCACCAGTAAGATGTGCAGGTGTAAACCCTGTGCCTTGTAACTGCAACCCTCCCGTGCCCACGGTATCTTCCAGAGGATGAGAACAAGCCGTAAGGAGATCACTTGAGGAAAAGAAAATGTTGGAAGAAGAGAGAAATCGGATCCTAACTAAACAGAAGCAAAGAATGAAAGAATTGCAGAAACTCCTGACAACCCGGGCTAAGGCTTATGACTCACATCAAAGTTTAGCT |
| --- | --- |
| *KIZ*:p.R76* | GGGCTGCGGGACAGTGAAAAGAAGAGATTGGACCTGGAAAAGAAACTTTATGAATATAATCAGTCTGATACATGCAGAGTTAAGCTGAAATATGTAAAACTAAAGAATTATCTGAAGGAAATATGTGAATCTGAAAAGAAGGCTCATACTTGAAACCAAGAATATTTAAAGCGATTTGAGCGTGTCCAAGCTCATGTTGTACACTTCACCACAAATACAGAGAAGCTTCAAAAACTGAAGCTCGAATATGAGACTCAAATTAAGAAGATGCTATGCTCAAAAGATAGCCTGGGACTAAAAGAG |
| *ABCA4*:p.R681* | GGAATCTACCTCCAGCAGATGCCCTACCCCTGCTTCGTGGACGATTCTTTCATGATCATCCTGAACCGCTGTTTCCCTATCTTCATGGTGCTGGCATGGATCTACTCTGTCTCCATGACTGTGAAGAGCATCGTCTTGGAGAAGGAGTTGTGACTGAAGGAGACCTTGAAAAATCAGGGTGTCTCCAATGCAGTGATTTGGTGTACCTGGTTCCTGGACAGCTTCTCCATCATGTCGATGAGCATCTTCCTCCTGACGATATTCATCATGCATGGAAGAATCCTACATTACAGCGACCCATTC |
| *ABCA4*:p.R20230* | ACCACATTCAAGATGCTCACTGGGGACACCACAGTGACCTCAGGGGATGCCACCGTAGCAGGCAAGAGTATTTTAACCAATATTTCTGAAGTCCATCAAAATATGGGCTACTGTCCTCAGTTTGATGCAATTGATGAGCTGCTCACAGGATGAGAACATCTTTACCTTTATGCCCGGCTTCGAGGTGTACCAGCAGAAGAAATCGAAAAGGTTGCAAACTGGAGTATTAAGAGCCTGGGCCTGACTGTCTACGCCGACTGCCTGGCTGGCACGTACAGTGGGGGCAACAAGCGGAAACTCTCC |
| *ABCA4*:p.R2149* | ACAGGGATGGACCCCCAGGCACGCCGCATGCTGTGGAACGTCATCGTGAGCATCATCAGAGAAGGGAGGGCTGTGGTCCTCACATCCCACAGCATGGAAGAATGTGAGGCACTGTGTACCCGGCTGGCCATCATGGTAAAGGGCGCCTTTTGATGTATGGGCACCATTCAGCATCTCAAGTCCAAATTTGGAGATGGCTATATCGTCACAATGAAGATCAAATCCCCGAAGGACGACCTGCTTCCTGACCTGAACCCTGTGGAGCAGTTCTTCCAGGGGAACTTCCCAGGCAGTGTGCAGAGG |
| *CERKL*:p.257* | GAATATGAAGGGCACGCTCTGTCACTGCTTAAGGAATGTGAACTCCAGGGATTTGATGGTGTTGTCTGTGTTGGTGGAGATGGATCTGCTAGCGAAGTAGCCCATGCTTTGCTTCTGAGAGCTCAGAAGAATGCTGGGATGGAAACAGACTGAATCCTGACTCCTGTCAGAGCACAGCTTCCACTTGGCTTAATACCAGCAGGATCTACCAATGTATTGGCACATTCTCTTCATGGAGTTCCTCATGTGATAACTGCAACATTGCACATTATAATGGGGCATGTACAGCTGGTCGACGTCTGC |
| *CNGA1*:p.R28* | AAGAACAATATTATCAATACACAGCAGTCTTTTGTAACCATGCCCAATGTGATTGTACCAGATATTGAAAAGGAAATATGAAGGATGGAAAATGGAGCATGCAGCTCCTTTTCTGAGGATGATGACAGTGCCTCTACATCTGAAGAATCAGAGAATGAAAAC |
| *PRCD*:p.R22* | TGCACCACCCTTTTCCTGCTCAGCACCCTGGCCATGCTCTGGCGCCGCCGATTTGCCAACTGAGTCCAACCAGAGCCCAGCGACGTGGATGGGGCAGCTAGGGGCAGCAGCTTGGATGCGGACCCTCAGTCCTCAGGCAGGGAGAAAGAACCTCTGAAG |
| *RP1*:p.R1993* | GATGCTATTAAAAACCAACCATTGCCTGGCAGTAATATGATTCATGGTACACTTCAGGAAGCTGACTCTTTGGATAAACTGTATGCTCTTTGTGGTCAACATTGCCCAATACTAACTGTTATTATCCAACCCATGAATGAGGAAGACTGAGGATTTGCATATCGCAAAGAATCTGATATTGAAAATTTCTTGGGTTTTTATTTATGGATGAAAATACACCCATATTTACTTCAGACAGACAAAAATGTGTTCAGGGAAGAGAACAATAAAGCAAGTATGAGACAAAATCTTATTGAT |
| *RP1*:p.R667* | TCTGAGGCTCCAGCTTCAGAAGCATCCTCTACTGTCACTGCAAGAATTGACAGACTAATTAATGAATTTGCTCAGTGTGGTTTAACAAAACTTCCAAAAAATGAAAAGAAGATTTTGTCATCTGTTGCCAGCAAAAAGAAGAAAAAATCTTGACAGCAAGCAATAAATTCCAGGTATCAAGATGGACAGCTTGCAACCAAAGGAATTCTTAATAAGAATGAGAGAATAAACACAAAAGGTAGAATTACAAAGGAAATGATAGTGCAAGATTCAGATAGTCCCCTTAAAGGAGGGATACTTTGT |
| *RP1L1*:p.R658* | AGGGAGCCTCTGGTTCTGGGCCTTTCCTGCTCCTGGGACTCGGAAGGAGCCTCTTCCACCCCTTCCACCTGCACTTCATCCCAGCAGGGGCAGAGAAGGCACAGAAGCCGGGCCAGTGCAATGTCCTCACCCAGCAGCCCTGGCCTTGGCTGAGTGGCCCCGAGAGGCCATCCCAGGCATTCTCACTACCGCAAGGACACCCACAGCCCACTGGACTCCTCTGTAACCAAGCAAGTGCCGAGGCCTCCTGAGCGGCGAAGGGCCTGCCAGGATGGCTCAGTGCCACGATATTCTGGAAGCTCA |
| *USH2A*:p.R737* | GATCCTGATGGCTGCAGTCCCTGTAACTGCAATACCTCTGGGACAGTGGATGGAGATATTACCTGTCACCAAAATTCAGGCCAGTGCAAGTGCAAAGCAAACGTTATTGGGCTTAGGTGTGATCATTGCAATTTTGGATTTAAATTTCTCTGAAGCTTTAATGATGTTGGATGTGAGCCCTGCCAGTGTAACCTCCATGGCTCAGTGAACAAATTCTGCAATCCTCACTCTGGGCAGTGTGAGTGCAAAAAAGAAGCCAAAGGACTTCAGTGTGACACCTGCAGAGAAAACTTTTATGGGTTA |
| *FAM161A*:p.R437* | GCCCAGGAGCATTTACAGAACTCATCTCCTCTGCCTTGTAGGTCAGCTTGTGGATGCAGGAACCCCAGGTGTCCTGAACAGGCTGTAAAGTTGAAGTGTAAACACAAGGTTAGGTGCCCAACTCCTGATTTTGAGGACCTTCCTGAGTGATACCAGAAACACCTCTCAGAACACAAGTCTCCAAAACTCTTAACAGTGTGTAAACCATTTGATCTTCATGCATCTCCACATGCATCTATTAAAAGAGAAAAAATTTTGGCAGACATCGAAGCAGATGAAGAAAATTTAAAAGAAACA |

**Supplementary Table 3: ACE-tRNA sequences used in this study**

| ACE-tRNA_V1 | GAGGGCCTATTTCCCATGATTCCTTCATATTTGCATATACGATACAAGGCTGTTAGAGAGATAATTGGAATTAATTTGACTGTAAACACAAAGATATTAGTACAAAATACGTGACGTAGAAAGTAATAATTTCTTGGGTAGTTTGCAGTTTTAAAATTATGTTTTAAAATGGACTATCATATGCTTACCGTAACTTGAAAGTATTTCGATTTCTTGGCTTTATATATCTTGTGGAAAGGACGAAACACCGGGCTCTGTGGCGCAATGGATAGCGCATTGGACTTCAAATTCAAAGGTTGTGGGTTCGAGTCCCACCAGAGTCGTTTTTT |
| --- | --- |
| ACE-tRNA_V2 | GAGGGCCTATTTCCCATGATTCCTTCATATTTGCATATACGATACAAGGCTGTTAGAGAGATAATTGGAATTAATTTGACTGTAAACACAAAGATATTAGTACAAAATACGTGACGTAGAAAGTAATAATTTCTTGGGTAGTTTGCAGTTTTAAAATTATGTTTTAAAATGGACTATCATATGCTTACCGTAACTTGAAAGTATTTCGATTTCTTGGCTTTATATATCTTGTGGAAAGGACGAAACACCGGGCTCTGTGGCGCAATGGATAGCGCATTGGACTTCAAATTCAAAGGTTGCGGGTTCGAGTCCCGTCAGAGTCGTTTTTT |
| ACE-tRNA_V3 | GAATTCCGGGACGGGGCGTTTTTTCCTTTAAAAAAAATTTAGAAAACCTCTTCCCGCGGGTGGCTCTGTGGCGCAATGGATAGCGCATTGGACTTCAAATTCAAAGGTTGCGGGTTCGAGTCCCGTCAGAGTCGGACCGTTTTTTTTTTTACGCCAAAACCAAAAAGAA |
| **Supplementary Table 4: Plasmid sequences used in this study** | |
| pDS-000 (Empty plasmid) | CTGCGCGCTCGCTCGCTCACTGAGGCCGCCCGGGCAAAGCCCGGGCGTCGGGCGACCTTTGGTCGCCCGGCCTCAGTGAGCGAGCGAGCGCGCAGAGAGGGAGTGGCCAACTCCATCACTAGGGGTTCCTTCTAGACAACTTTGTATAGAAAAGTTGAAGCTTGGTACCACCGGCCGCTCGAGGTTTAAACCTCGACATTGATTATTGACTAGTTATTAATAGTAATCAATTACGGGGTCATTAGTTCATAGCCCATATATGGAGTTCCGCGTTACATAACTTACGGTAAATGGCCCGCCTGGCTGACCGCCCAACGACCCCCGCCCATTGACGTCAATAATGACGTATGTTCCCATAGTAACGCCAATAGGGACTTTCCATTGACGTCAATGGGTGGAGTATTTACGGTAAACTGCCCACTTGGCAGTACATCAAGTGTATCATATGCCAAGTACGCCCCCTATTGACGTCAATGACGGTAAATGGCCCGCCTGGCATTATGCCCAGTACATGACCTTATGGGACTTTCCTACTTGGCAGTACATCTACGTATTAGTCATCGCTATTACCATGGTCGAGGTGAGCCCCACGTTCTGCTTCACTCTCCCCATCTCCCCCCCCTCCCCACCCCCAATTTTGTATTTATTTATTTTTTAATTATTTTGTGCAGCGATGGGGGCGGGGGGGGGGGGGGGGCGCGCGCCAGGCGGGGCGGGGCGGGGCGAGGGGCGGGGCGGGGCGAGGCGGAGAGGTGCGGCGGCAGCCAATCAGAGCGGCGCGCTCCGAAAGTTTCCTTTTATGGCGAGGCGGCGGCGGCGGCGGCCCTATAAAAAGCGAAGCGCGCGGCGGGCGGGAGTCGCTGCGCGCTGCCTTCGCCCCGTGCCCCGCTCCGCCGCCGCCTCGCGCCGCCCGCCCCGGCTCTGACTGACCGCGTTACTCCCACAGGTGAGCGGGCGGGACGGCCCTTCTCCTCCGGGCTGTAATTAGCGCTTGGTTTAATGACGGCTTGTTTCTTTTCTGTGGCTGCGTGAAAGCCTTGAGGGGCTCCGGGAGGGCCCTTTGTGCGGGGGGAGCGGCTCGGGGGGTGCGTGCGTGTGTGTGTGCGTGGGGAGCGCCGCGTGCGGCTCCGCGCTGCCCGGCGGCTGTGAGCGCTGCGGGCGCGGCGCGGGGCTTTGTGCGCTCCGCAGTGTGCGCGAGGGGAGCGCGGCCGGGGGCGGTGCCCCGCGGTGCGGGGGGGGCTGCGAGGGGAACAAAGGCTGCGTGCGGGGTGTGTGCGTGGGGGGGTGAGCAGGGGGTGTGGGCGCGTCGGTCGGGCTGCAACCCCCCCTGCACCCCCCTCCCCGAGTTGCTGAGCACGGCCCGGCTTCGGGTGCGGGGCTCCGTACGGGGCGTGGCGCGGGGCTCGCCGTGCCGGGCGGGGGGTGGCGGCAGGTGGGGGTGCCGGGCGGGGCGGGGCCGCCTCGGGCCGGGGAGGGCTCGGGGGAGGGGCGCGGCGGCCCCCGGAGCGCCGGCGGCTGTCGAGGCGCGGCGAGCCGCAGCCATTGCCTTTTATGGTAATCGTGCGAGAGGGCGCAGGGACTTCCTTTGTCCCAAATCTGTGCGGAGCCGAAATCTGGGAGGCGCCGCCGCACCCCCTCTAGCGGGCGCGGGGCGAAGCGGTGCGGCGCCGGCAGGAAGGAAATGGGCGGGGAGGGCCTTCGTGCGTCGCCGCGCCGCCGTCCCCTTCTCCCTCTCCAGCCTCGGGGCTGTCCGCGGGGGGACGGCTGCCTTCGGGGGGGACGGGGCAGGGCGGGGTTCGGCTTCTGGCGTGTGACCGGCGGCTCTAGAGCCTCTGCTAACCATGTTCATGCCTTCTTCTTTTTCCTACAGCTCCTGGGCAACGTGCTGGTTATTGTGCTGTCTCATCATTTTGGCAAAGAATTGACCGGTAGATCTACATCGATGGACGCGTTTAATTAAGTCGACCAAGTTTGTACAAAAAAGCAGGCTGCCACCATGGTGAGCAAGGGCGAGGAGGATAACATGGCCATCATCAAGGAGTTCATGCGCTTCAAGGTGCACATGGAGGGCTCCGTGAACGGCCACGAGTTCGAGATCGAGGGCGAGGGCGAGGGCCGCCCCTACGAGGGCACCCAGACCGCCAAGCTGAAGGTGACCAAGGGTGGCCCCCTGCCCTTCGCCTGGGACATCCTGTCCCCTCAGTTCATGTACGGCTCCAAGGCCTACGTGAAGCACCCCGCCGACATCCCCGACTACTTGAAGCTGTCCTTCCCCGAGGGCTTCAAGTGGGAGCGCGTGATGAACTTCGAGGACGGCGGCGTGGTGACCGTGACCCAGGACTCCTCCCTGCAGGACGGCGAGTTCATCTACAAGGTGAAGCTGCGCGGCACCAACTTCCCCTCCGACGGCCCCGTAATGCAGAAGAAGACCATGGGCTGGGAGGCCTCCTCCGAGCGGATGTACCCCGAGGACGGCGCCCTGAAGGGCGAGATCAAGCAGAGGCTGAAGCTGAAGGACGGCGGCCACTACGACGCTGAGGTCAAGACCACCTACAAGGCCAAGAAGCCCGTGCAGCTGCCCGGCGCCTACAACGTCAACATCAAGTTGGACATCACCTCCCACAACGAGGACTACACCATCGTGGAACAGTACGAACGCGCCGAGGGCCGCCACTCCACCGGCGGCATGGACGAGCTGTACAAGTAAACCCAGCTTTCTTGTACAAAGTGGGAATTCCGATAATCAACCTCTGGATTACAAAATTTGTGAAAGATTGACTGGTATTCTTAACTATGTTGCTCCTTTTACGCTATGTGGATACGCTGCTTTAATGCCTTTGTATCATGCTATTGCTTCCCGTATGGCTTTCATTTTCTCCTCCTTGTATAAATCCTGGTTGCTGTCTCTTTATGAGGAGTTGTGGCCCGTTGTCAGGCAACGTGGCGTGGTGTGCACTGTGTTTGCTGACGCAACCCCCACTGGTTGGGGCATTGCCACCACCTGTCAGCTCCTTTCCGGGACTTTCGCTTTCCCCCTCCCTATTGCCACGGCGGAACTCATCGCCGCCTGCCTTGCCCGCTGCTGGACAGGGGCTCGGCTGTTGGGCACTGACAATTCCGTGGTGTTGTCGGGGAAGCTGACGTCCTTTCCATGGCTGCTCGCCTGTGTTGCCACCTGGATTCTGCGCGGGACGTCCTTCTGCTACGTCCCTTCGGCCCTCAATCCAGCGGACCTTCCTTCCCGCGGCCTGCTGCCGGCTCTGCGGCCTCTTCCGCGTCTTCGCCTTCGCCCTCAGACGAGTCGGATCTCCCTTTGGGCCGCCTCCCCGCATCGGGAATTCCTAGAGCTCGCTGATCAGCCTCGACTGTGCCTTCTAGTTGCCAGCCATCTGTTGTTTGCCCCTCCCCCGTGCCTTCCTTGACCCTGGAAGGTGCCACTCCCACTGTCCTTTCCTAATAAAATGAGGAAATTGCATCGCATTGTCTGAGTAGGTGTCATTCTATTCTGGGGGGTGGGGTGGGGCAGGACAGCAAGGGGGAGGATTGGGAAGAGAATAGCAGGCATGCTGGGGAGGGCCGCAGGAACCCCTAGTGATGGAGTTGGCCACTCCCTCTCTGCGCGCTCGCTCGCTCACTGAGGCCGGGCGACCAAAGGTCGCCCGACGCCCGGGCTTTGCCCGGGCGGCCTCAGTGAGCGAGCGAGCGCGCAGCTGCCTGCAGGGGCGCCTGATGCGGTATTTTCTCCTTACGCATCTGTGCGGTATTTCACACCGCATACGTCAAAGCAACCATAGTACGCGCCCTGTAGCGGCGCATTAAGCGCGGCGGGTGTGGTGGTTACGCGCAGCGTGACCGCTACACTTGCCAGCGCCTTAGCGCCCGCTCCTTTCGCTTTCTTCCCTTCCTTTCTCGCCACGTTCGCCGGCTTTCCCCGTCAAGCTCTAAATCGGGGGCTCCCTTTAGGGTTCCGATTTAGTGCTTTACGGCACCTCGACCCCAAAAAACTTGATTTGGGTGATGGTTCACGTAGTGGGCCATCGCCCTGATAGACGGTTTTTCGCCCTTTGACGTTGGAGTCCACGTTCTTTAATAGTGGACTCTTGTTCCAAACTGGAACAACACTCAACTCTATCTCGGGCTATTCTTTTGATTTATAAGGGATTTTGCCGATTTCGGTCTATTGGTTAAAAAATGAGCTGATTTAACAAAAATTTAACGCGAATTTTAACAAAATATTAACGTTTACAATTTTATGGTGCACTCTCAGTACAATCTGCTCTGATGCCGCATAGTTAAGCCAGCCCCGACACCCGCCAACACCCGCTGACGCGCCCTGACGGGCTTGTCTGCTCCCGGCATCCGCTTACAGACAAGCTGTGACCGTCTCCGGGAGCTGCATGTGTCAGAGGTTTTCACCGTCATCACCGAAACGCGCGAGACGAAAGGGCCTCGTGATACGCCTATTTTTATAGGTTAATGTCATGATAATAATGGTTTCTTAGACGTCAGGTGGCACTTTTCGGGGAAATGTGCGCGGAACCCCTATTTGTTTATTTTTCTAAATACATTCAAATATGTATCCGCTCATGAGACAATAACCCTGATAAATGCTTCAATAATATTGAAAAAGGAAGAGTATGAGTATTCAACATTTCCGTGTCGCCCTTATTCCCTTTTTTGCGGCATTTTGCCTTCCTGTTTTTGCTCACCCAGAAACGCTGGTGAAAGTAAAAGATGCTGAAGATCAGTTGGGTGCACGAGTGGGTTACATCGAACTGGATCTCAACAGCGGTAAGATCCTTGAGAGTTTTCGCCCCGAAGAACGTTTTCCAATGATGAGCACTTTTAAAGTTCTGCTATGTGGCGCGGTATTATCCCGTATTGACGCCGGGCAAGAGCAACTCGGTCGCCGCATACACTATTCTCAGAATGACTTGGTTGAGTACTCACCAGTCACAGAAAAGCATCTTACGGATGGCATGACAGTAAGAGAATTATGCAGTGCTGCCATAACCATGAGTGATAACACTGCGGCCAACTTACTTCTGACAACGATCGGAGGACCGAAGGAGCTAACCGCTTTTTTGCACAACATGGGGGATCATGTAACTCGCCTTGATCGTTGGGAACCGGAGCTGAATGAAGCCATACCAAACGACGAGCGTGACACCACGATGCCTGTAGCAATGGCAACAACGTTGCGCAAACTATTAACTGGCGAACTACTTACTCTAGCTTCCCGGCAACAATTAATAGACTGGATGGAGGCGGATAAAGTTGCAGGACCACTTCTGCGCTCGGCCCTTCCGGCTGGCTGGTTTATTGCTGATAAATCTGGAGCCGGTGAGCGTGGAAGCCGCGGTATCATTGCAGCACTGGGGCCAGATGGTAAGCCCTCCCGTATCGTAGTTATCTACACGACGGGGAGTCAGGCAACTATGGATGAACGAAATAGACAGATCGCTGAGATAGGTGCCTCACTGATTAAGCATTGGTAACTGTCAGACCAAGTTTACTCATATATACTTTAGATTGATTTAAAACTTCATTTTTAATTTAAAAGGATCTAGGTGAAGATCCTTTTTGATAATCTCATGACCAAAATCCCTTAACGTGAGTTTTCGTTCCACTGAGCGTCAGACCCCGTAGAAAAGATCAAAGGATCTTCTTGAGATCCTTTTTTTCTGCGCGTAATCTGCTGCTTGCAAACAAAAAAACCACCGCTACCAGCGGTGGTTTGTTTGCCGGATCAAGAGCTACCAACTCTTTTTCCGAAGGTAACTGGCTTCAGCAGAGCGCAGATACCAAATACTGTTCTTCTAGTGTAGCCGTAGTTAGGCCACCACTTCAAGAACTCTGTAGCACCGCCTACATACCTCGCTCTGCTAATCCTGTTACCAGTGGCTGCTGCCAGTGGCGATAAGTCGTGTCTTACCGGGTTGGACTCAAGACGATAGTTACCGGATAAGGCGCAGCGGTCGGGCTGAACGGGGGGTTCGTGCACACAGCCCAGCTTGGAGCGAACGACCTACACCGAACTGAGATACCTACAGCGTGAGCTATGAGAAAGCGCCACGCTTCCCGAAGGGAGAAAGGCGGACAGGTATCCGGTAAGCGGCAGGGTCGGAACAGGAGAGCGCACGAGGGAGCTTCCAGGGGGAAACGCCTGGTATCTTTATAGTCCTGTCGGGTTTCGCCACCTCTGACTTGAGCGTCGATTTTTGTGATGCTCGTCAGGGGGGCGGAGCCTATGGAAAAACGCCAGCAACGCGGCCTTTTTACGGTTCCTGGCCTTTTGCTGGCCTTTTGCTCACATGTCCTGCAGGCAG |
| pDS-001 (With stuffer sequence) | CTGCGCGCTCGCTCGCTCACTGAGGCCGCCCGGGCAAAGCCCGGGCGTCGGGCGACCTTTGGTCGCCCGGCCTCAGTGAGCGAGCGAGCGCGCAGAGAGGGAGTGGCCAACTCCATCACTAGGGGTTCCTTCTAGACAACTTTGTATAGAAAAGTTGAAGCTTGGTACCACCGGCCGCTCGAGGAATTCCGGGACGGGGCGTTTTTTCCTTTAAAAAAAATTTAGAAAACCTCTTCCCGCGGGTGGCTCTGTGGCGCAATGGATAGCGCATTGGACTTCAAATTCAAAGGTTGCGGGTTCGAGTCCCGTCAGAGTCGGACCGTTTTTTTTTTTACGCCAAAACCAAAAAGAAACTAGTCATATGGGGCCCACGCGTGTCGACACCGTTctccctcgaatttgaaagagatatttgctgcccggtacttccatgtaacaacgcttgttacgaagaagtgttcgctaaacgtcccagtggtgcccccgatcgtaatgtgcgccttaatgctagatagacatctgtctgctgaaattaaagacacgtcgcacctcttgctagcaatcctcgtggcgctacgctctctgaaattcccgtgattctcgattagcataatgcctatacccttgctattaagctatatcgcattagatcagcgataatgatgaagaccagacgcgcgcctgttgtgtattgaggcctggcggaatagataccttcattcgtcgggtacccttttttgcatcggcgcaaatggatagtttagatgccaaagttttacctggcgactgacgcgagtgaatcctctcagttggataggtttacgaggtctgcgatcagaagaacagtcaaaaccgaatgattacgtcccgctactaaccgagcccctttcgctgggcggtacacccgcactgtgcgctgaaacccggtatgcgcggggcctaaccagcagcgttacgagagcccacaaccaaggtagcccggcgtttccgagcgaatcagtgggtctcgtacactccttgcggttttcttggacatgcaaagaacgcgaaagcaacggtggcgctcgggcccggccagaatagcgcttataagacgtacaagagcagcatcaagccataagacgcaaaggtgctaggtcaagctcacgatttgtgcgactatatgctatccgaaccgtgaaagtagttgccttcggactcagtcaatggttgtcccctccatgatagggtctcacctctggcgggggtcgagtccggtgcgcttgcatcgttcgattagtgacgacaatgccgctactgcacgtccgcacgaccgcctaactgagacccatttggttgtgtcgggtttcgacgggggaccatgcctgtagatgccactaatagcgtcctatgtgacgcgttgtgctgtccgcacaatctcctcaaattcctagctcgcgaataaatgtcaaaagccaggatctccccctgctgcgggacccggggatctagggcttcaccgcatgtgcaggcgattgttagatgcgaagggttttactctattcccacggagctcatgcacctccgaggtcggacgcatcccgtattaagccctgtatgggtagtctaggatcttaagaacccggcataggttcagatcagttacccatcagagagaaatcccaggcagagattgtcctacttcggtaacattaggacagaccctggccgcgcgtaggcctatctgcacccccgcgtttaatccgtcaagggtcctcacagccgctcgtggacctctgcgggactccaaatgtcagtggtgaactgccgtacactgccattgaatttggcgtacaccgcccgaactaggggctcgccgcacacaaagtcgacaaactatcctcagtcgtacaaggggcgataaggtagcagacttctaccggagggatttatgatatatcagtggtgcctcctaaggagtcgtcagtcctgctgacctgttggacgactatcaaaattcaccaaggagcagttccgtcaaaacggcaaggcgtcttcccaatcatcgggaattgctggtcttgaaccagtaaccacatgctgccgtacagctagctgaagctgctaatcgttatcgctatcattccgaccaagcaagtatatcttctggtcttgatcgatgttgcgttaatcgtcattacaggtgtgagcatgatcaactttcggttatttttgcgttgtcggacccggcaccgcttcacctccgttggcatgaacccaccggggatttggaagctgtcagagacaagatcttctagagattaggatcggtctggggcgttgaagcacctacgaaagtgctatctcccgacttatcactacgattgctcctctctcaattttaataacatagtgtgggggGCATGCTGGGGAGGGCCGCAGGAACCCCTAGTGATGGAGTTGGCCACTCCCTCTCTGCGCGCTCGCTCGCTCACTGAGGCCGGGCGACCAAAGGTCGCCCGACGCCCGGGCTTTGCCCGGGCGGCCTCAGTGAGCGAGCGAGCGCGCAGCTGCCTGCAGGGGCGCCTGATGCGGTATTTTCTCCTTACGCATCTGTGCGGTATTTCACACCGCATACGTCAAAGCAACCATAGTACGCGCCCTGTAGCGGCGCATTAAGCGCGGCGGGTGTGGTGGTTACGCGCAGCGTGACCGCTACACTTGCCAGCGCCTTAGCGCCCGCTCCTTTCGCTTTCTTCCCTTCCTTTCTCGCCACGTTCGCCGGCTTTCCCCGTCAAGCTCTAAATCGGGGGCTCCCTTTAGGGTTCCGATTTAGTGCTTTACGGCACCTCGACCCCAAAAAACTTGATTTGGGTGATGGTTCACGTAGTGGGCCATCGCCCTGATAGACGGTTTTTCGCCCTTTGACGTTGGAGTCCACGTTCTTTAATAGTGGACTCTTGTTCCAAACTGGAACAACACTCAACTCTATCTCGGGCTATTCTTTTGATTTATAAGGGATTTTGCCGATTTCGGTCTATTGGTTAAAAAATGAGCTGATTTAACAAAAATTTAACGCGAATTTTAACAAAATATTAACGTTTACAATTTTATGGTGCACTCTCAGTACAATCTGCTCTGATGCCGCATAGTTAAGCCAGCCCCGACACCCGCCAACACCCGCTGACGCGCCCTGACGGGCTTGTCTGCTCCCGGCATCCGCTTACAGACAAGCTGTGACCGTCTCCGGGAGCTGCATGTGTCAGAGGTTTTCACCGTCATCACCGAAACGCGCGAGACGAAAGGGCCTCGTGATACGCCTATTTTTATAGGTTAATGTCATGATAATAATGGTTTCTTAGACGTCAGGTGGCACTTTTCGGGGAAATGTGCGCGGAACCCCTATTTGTTTATTTTTCTAAATACATTCAAATATGTATCCGCTCATGAGACAATAACCCTGATAAATGCTTCAATAATATTGAAAAAGGAAGAGTATGAGTATTCAACATTTCCGTGTCGCCCTTATTCCCTTTTTTGCGGCATTTTGCCTTCCTGTTTTTGCTCACCCAGAAACGCTGGTGAAAGTAAAAGATGCTGAAGATCAGTTGGGTGCACGAGTGGGTTACATCGAACTGGATCTCAACAGCGGTAAGATCCTTGAGAGTTTTCGCCCCGAAGAACGTTTTCCAATGATGAGCACTTTTAAAGTTCTGCTATGTGGCGCGGTATTATCCCGTATTGACGCCGGGCAAGAGCAACTCGGTCGCCGCATACACTATTCTCAGAATGACTTGGTTGAGTACTCACCAGTCACAGAAAAGCATCTTACGGATGGCATGACAGTAAGAGAATTATGCAGTGCTGCCATAACCATGAGTGATAACACTGCGGCCAACTTACTTCTGACAACGATCGGAGGACCGAAGGAGCTAACCGCTTTTTTGCACAACATGGGGGATCATGTAACTCGCCTTGATCGTTGGGAACCGGAGCTGAATGAAGCCATACCAAACGACGAGCGTGACACCACGATGCCTGTAGCAATGGCAACAACGTTGCGCAAACTATTAACTGGCGAACTACTTACTCTAGCTTCCCGGCAACAATTAATAGACTGGATGGAGGCGGATAAAGTTGCAGGACCACTTCTGCGCTCGGCCCTTCCGGCTGGCTGGTTTATTGCTGATAAATCTGGAGCCGGTGAGCGTGGAAGCCGCGGTATCATTGCAGCACTGGGGCCAGATGGTAAGCCCTCCCGTATCGTAGTTATCTACACGACGGGGAGTCAGGCAACTATGGATGAACGAAATAGACAGATCGCTGAGATAGGTGCCTCACTGATTAAGCATTGGTAACTGTCAGACCAAGTTTACTCATATATACTTTAGATTGATTTAAAACTTCATTTTTAATTTAAAAGGATCTAGGTGAAGATCCTTTTTGATAATCTCATGACCAAAATCCCTTAACGTGAGTTTTCGTTCCACTGAGCGTCAGACCCCGTAGAAAAGATCAAAGGATCTTCTTGAGATCCTTTTTTTCTGCGCGTAATCTGCTGCTTGCAAACAAAAAAACCACCGCTACCAGCGGTGGTTTGTTTGCCGGATCAAGAGCTACCAACTCTTTTTCCGAAGGTAACTGGCTTCAGCAGAGCGCAGATACCAAATACTGTTCTTCTAGTGTAGCCGTAGTTAGGCCACCACTTCAAGAACTCTGTAGCACCGCCTACATACCTCGCTCTGCTAATCCTGTTACCAGTGGCTGCTGCCAGTGGCGATAAGTCGTGTCTTACCGGGTTGGACTCAAGACGATAGTTACCGGATAAGGCGCAGCGGTCGGGCTGAACGGGGGGTTCGTGCACACAGCCCAGCTTGGAGCGAACGACCTACACCGAACTGAGATACCTACAGCGTGAGCTATGAGAAAGCGCCACGCTTCCCGAAGGGAGAAAGGCGGACAGGTATCCGGTAAGCGGCAGGGTCGGAACAGGAGAGCGCACGAGGGAGCTTCCAGGGGGAAACGCCTGGTATCTTTATAGTCCTGTCGGGTTTCGCCACCTCTGACTTGAGCGTCGATTTTTGTGATGCTCGTCAGGGGGGCGGAGCCTTGGAAAAACGCCAGCAACGCGGCCTTTTTACGGTTCCTGGCCTTTTGCTGGCCTTTTGCTCACATGTCCTGCAGGCAG |
| **pDS-002 (With mCherry)** | CTGCGCGCTCGCTCGCTCACTGAGGCCGCCCGGGCAAAGCCCGGGCGTCGGGCGACCTTTGGTCGCCCGGCCTCAGTGAGCGAGCGAGCGCGCAGAGAGGGAGTGGCCAACTCCATCACTAGGGGTTCCTTCTAGACAACTTTGTATAGAAAAGTTGAAGCTTGGTACCGAATTCCGGGACGGGGCGTTTTTTCCTTTAAAAAAAATTTAGAAAACCTCTTCCCGCGGGTGGCTCTGTGGCGCAATGGATAGCGCATTGGACTTCAAATTCAAAGGTTGCGGGTTCGAGTCCCGTCAGAGTCGGACCGTTTTTTTTTTTACGCCAAAACCAAAAAGAACTCGAGGTTTAAACCTCGACATTGATTATTGACTAGTTATTAATAGTAATCAATTACGGGGTCATTAGTTCATAGCCCATATATGGAGTTCCGCGTTACATAACTTACGGTAAATGGCCCGCCTGGCTGACCGCCCAACGACCCCCGCCCATTGACGTCAATAATGACGTATGTTCCCATAGTAACGCCAATAGGGACTTTCCATTGACGTCAATGGGTGGAGTATTTACGGTAAACTGCCCACTTGGCAGTACATCAAGTGTATCATATGCCAAGTACGCCCCCTATTGACGTCAATGACGGTAAATGGCCCGCCTGGCATTATGCCCAGTACATGACCTTATGGGACTTTCCTACTTGGCAGTACATCTACGTATTAGTCATCGCTATTACCATGGTCGAGGTGAGCCCCACGTTCTGCTTCACTCTCCCCATCTCCCCCCCCTCCCCACCCCCAATTTTGTATTTATTTATTTTTTAATTATTTTGTGCAGCGATGGGGGCGGGGGGGGGGGGGGGGCGCGCGCCAGGCGGGGCGGGGCGGGGCGAGGGGCGGGGCGGGGCGAGGCGGAGAGGTGCGGCGGCAGCCAATCAGAGCGGCGCGCTCCGAAAGTTTCCTTTTATGGCGAGGCGGCGGCGGCGGCGGCCCTATAAAAAGCGAAGCGCGCGGCGGGCGGGAGTCGCTGCGCGCTGCCTTCGCCCCGTGCCCCGCTCCGCCGCCGCCTCGCGCCGCCCGCCCCGGCTCTGACTGACCGCGTTACTCCCACAGGTGAGCGGGCGGGACGGCCCTTCTCCTCCGGGCTGTAATTAGCGCTTGGTTTAATGACGGCTTGTTTCTTTTCTGTGGCTGCGTGAAAGCCTTGAGGGGCTCCGGGAGGGCCCTTTGTGCGGGGGGAGCGGCTCGGGGGGTGCGTGCGTGTGTGTGTGCGTGGGGAGCGCCGCGTGCGGCTCCGCGCTGCCCGGCGGCTGTGAGCGCTGCGGGCGCGGCGCGGGGCTTTGTGCGCTCCGCAGTGTGCGCGAGGGGAGCGCGGCCGGGGGCGGTGCCCCGCGGTGCGGGGGGGGCTGCGAGGGGAACAAAGGCTGCGTGCGGGGTGTGTGCGTGGGGGGGTGAGCAGGGGGTGTGGGCGCGTCGGTCGGGCTGCAACCCCCCCTGCACCCCCCTCCCCGAGTTGCTGAGCACGGCCCGGCTTCGGGTGCGGGGCTCCGTACGGGGCGTGGCGCGGGGCTCGCCGTGCCGGGCGGGGGGTGGCGGCAGGTGGGGGTGCCGGGCGGGGCGGGGCCGCCTCGGGCCGGGGAGGGCTCGGGGGAGGGGCGCGGCGGCCCCCGGAGCGCCGGCGGCTGTCGAGGCGCGGCGAGCCGCAGCCATTGCCTTTTATGGTAATCGTGCGAGAGGGCGCAGGGACTTCCTTTGTCCCAAATCTGTGCGGAGCCGAAATCTGGGAGGCGCCGCCGCACCCCCTCTAGCGGGCGCGGGGCGAAGCGGTGCGGCGCCGGCAGGAAGGAAATGGGCGGGGAGGGCCTTCGTGCGTCGCCGCGCCGCCGTCCCCTTCTCCCTCTCCAGCCTCGGGGCTGTCCGCGGGGGGACGGCTGCCTTCGGGGGGGACGGGGCAGGGCGGGGTTCGGCTTCTGGCGTGTGACCGGCGGCTCTAGAGCCTCTGCTAACCATGTTCATGCCTTCTTCTTTTTCCTACAGCTCCTGGGCAACGTGCTGGTTATTGTGCTGTCTCATCATTTTGGCAAAGAATTGACCGGTAGATCTACATCGATGGACGCGTTTAATTAAGTCGACCAAGTTTGTACAAAAAAGCAGGCTGCCACCATGGTGAGCAAGGGCGAGGAGGATAACATGGCCATCATCAAGGAGTTCATGCGCTTCAAGGTGCACATGGAGGGCTCCGTGAACGGCCACGAGTTCGAGATCGAGGGCGAGGGCGAGGGCCGCCCCTACGAGGGCACCCAGACCGCCAAGCTGAAGGTGACCAAGGGTGGCCCCCTGCCCTTCGCCTGGGACATCCTGTCCCCTCAGTTCATGTACGGCTCCAAGGCCTACGTGAAGCACCCCGCCGACATCCCCGACTACTTGAAGCTGTCCTTCCCCGAGGGCTTCAAGTGGGAGCGCGTGATGAACTTCGAGGACGGCGGCGTGGTGACCGTGACCCAGGACTCCTCCCTGCAGGACGGCGAGTTCATCTACAAGGTGAAGCTGCGCGGCACCAACTTCCCCTCCGACGGCCCCGTAATGCAGAAGAAGACCATGGGCTGGGAGGCCTCCTCCGAGCGGATGTACCCCGAGGACGGCGCCCTGAAGGGCGAGATCAAGCAGAGGCTGAAGCTGAAGGACGGCGGCCACTACGACGCTGAGGTCAAGACCACCTACAAGGCCAAGAAGCCCGTGCAGCTGCCCGGCGCCTACAACGTCAACATCAAGTTGGACATCACCTCCCACAACGAGGACTACACCATCGTGGAACAGTACGAACGCGCCGAGGGCCGCCACTCCACCGGCGGCATGGACGAGCTGTACAAGTAAACCCAGCTTTCTTGTACAAAGTGGGAATTCCGATAATCAACCTCTGGATTACAAAATTTGTGAAAGATTGACTGGTATTCTTAACTATGTTGCTCCTTTTACGCTATGTGGATACGCTGCTTTAATGCCTTTGTATCATGCTATTGCTTCCCGTATGGCTTTCATTTTCTCCTCCTTGTATAAATCCTGGTTGCTGTCTCTTTATGAGGAGTTGTGGCCCGTTGTCAGGCAACGTGGCGTGGTGTGCACTGTGTTTGCTGACGCAACCCCCACTGGTTGGGGCATTGCCACCACCTGTCAGCTCCTTTCCGGGACTTTCGCTTTCCCCCTCCCTATTGCCACGGCGGAACTCATCGCCGCCTGCCTTGCCCGCTGCTGGACAGGGGCTCGGCTGTTGGGCACTGACAATTCCGTGGTGTTGTCGGGGAAGCTGACGTCCTTTCCATGGCTGCTCGCCTGTGTTGCCACCTGGATTCTGCGCGGGACGTCCTTCTGCTACGTCCCTTCGGCCCTCAATCCAGCGGACCTTCCTTCCCGCGGCCTGCTGCCGGCTCTGCGGCCTCTTCCGCGTCTTCGCCTTCGCCCTCAGACGAGTCGGATCTCCCTTTGGGCCGCCTCCCCGCATCGGGAATTCCTAGAGCTCGCTGATCAGCCTCGACTGTGCCTTCTAGTTGCCAGCCATCTGTTGTTTGCCCCTCCCCCGTGCCTTCCTTGACCCTGGAAGGTGCCACTCCCACTGTCCTTTCCTAATAAAATGAGGAAATTGCATCGCATTGTCTGAGTAGGTGTCATTCTATTCTGGGGGGTGGGGTGGGGCAGGACAGCAAGGGGGAGGATTGGGAAGAGAATAGCAGGCATGCTGGGGAGGGCCGCAGGAACCCCTAGTGATGGAGTTGGCCACTCCCTCTCTGCGCGCTCGCTCGCTCACTGAGGCCGGGCGACCAAAGGTCGCCCGACGCCCGGGCTTTGCCCGGGCGGCCTCAGTGAGCGAGCGAGCGCGCAGCTGCCTGCAGGGGCGCCTGATGCGGTATTTTCTCCTTACGCATCTGTGCGGTATTTCACACCGCATACGTCAAAGCAACCATAGTACGCGCCCTGTAGCGGCGCATTAAGCGCGGCGGGTGTGGTGGTTACGCGCAGCGTGACCGCTACACTTGCCAGCGCCTTAGCGCCCGCTCCTTTCGCTTTCTTCCCTTCCTTTCTCGCCACGTTCGCCGGCTTTCCCCGTCAAGCTCTAAATCGGGGGCTCCCTTTAGGGTTCCGATTTAGTGCTTTACGGCACCTCGACCCCAAAAAACTTGATTTGGGTGATGGTTCACGTAGTGGGCCATCGCCCTGATAGACGGTTTTTCGCCCTTTGACGTTGGAGTCCACGTTCTTTAATAGTGGACTCTTGTTCCAAACTGGAACAACACTCAACTCTATCTCGGGCTATTCTTTTGATTTATAAGGGATTTTGCCGATTTCGGTCTATTGGTTAAAAAATGAGCTGATTTAACAAAAATTTAACGCGAATTTTAACAAAATATTAACGTTTACAATTTTATGGTGCACTCTCAGTACAATCTGCTCTGATGCCGCATAGTTAAGCCAGCCCCGACACCCGCCAACACCCGCTGACGCGCCCTGACGGGCTTGTCTGCTCCCGGCATCCGCTTACAGACAAGCTGTGACCGTCTCCGGGAGCTGCATGTGTCAGAGGTTTTCACCGTCATCACCGAAACGCGCGAGACGAAAGGGCCTCGTGATACGCCTATTTTTATAGGTTAATGTCATGATAATAATGGTTTCTTAGACGTCAGGTGGCACTTTTCGGGGAAATGTGCGCGGAACCCCTATTTGTTTATTTTTCTAAATACATTCAAATATGTATCCGCTCATGAGACAATAACCCTGATAAATGCTTCAATAATATTGAAAAAGGAAGAGTATGAGTATTCAACATTTCCGTGTCGCCCTTATTCCCTTTTTTGCGGCATTTTGCCTTCCTGTTTTTGCTCACCCAGAAACGCTGGTGAAAGTAAAAGATGCTGAAGATCAGTTGGGTGCACGAGTGGGTTACATCGAACTGGATCTCAACAGCGGTAAGATCCTTGAGAGTTTTCGCCCCGAAGAACGTTTTCCAATGATGAGCACTTTTAAAGTTCTGCTATGTGGCGCGGTATTATCCCGTATTGACGCCGGGCAAGAGCAACTCGGTCGCCGCATACACTATTCTCAGAATGACTTGGTTGAGTACTCACCAGTCACAGAAAAGCATCTTACGGATGGCATGACAGTAAGAGAATTATGCAGTGCTGCCATAACCATGAGTGATAACACTGCGGCCAACTTACTTCTGACAACGATCGGAGGACCGAAGGAGCTAACCGCTTTTTTGCACAACATGGGGGATCATGTAACTCGCCTTGATCGTTGGGAACCGGAGCTGAATGAAGCCATACCAAACGACGAGCGTGACACCACGATGCCTGTAGCAATGGCAACAACGTTGCGCAAACTATTAACTGGCGAACTACTTACTCTAGCTTCCCGGCAACAATTAATAGACTGGATGGAGGCGGATAAAGTTGCAGGACCACTTCTGCGCTCGGCCCTTCCGGCTGGCTGGTTTATTGCTGATAAATCTGGAGCCGGTGAGCGTGGAAGCCGCGGTATCATTGCAGCACTGGGGCCAGATGGTAAGCCCTCCCGTATCGTAGTTATCTACACGACGGGGAGTCAGGCAACTATGGATGAACGAAATAGACAGATCGCTGAGATAGGTGCCTCACTGATTAAGCATTGGTAACTGTCAGACCAAGTTTACTCATATATACTTTAGATTGATTTAAAACTTCATTTTTAATTTAAAAGGATCTAGGTGAAGATCCTTTTTGATAATCTCATGACCAAAATCCCTTAACGTGAGTTTTCGTTCCACTGAGCGTCAGACCCCGTAGAAAAGATCAAAGGATCTTCTTGAGATCCTTTTTTTCTGCGCGTAATCTGCTGCTTGCAAACAAAAAAACCACCGCTACCAGCGGTGGTTTGTTTGCCGGATCAAGAGCTACCAACTCTTTTTCCGAAGGTAACTGGCTTCAGCAGAGCGCAGATACCAAATACTGTTCTTCTAGTGTAGCCGTAGTTAGGCCACCACTTCAAGAACTCTGTAGCACCGCCTACATACCTCGCTCTGCTAATCCTGTTACCAGTGGCTGCTGCCAGTGGCGATAAGTCGTGTCTTACCGGGTTGGACTCAAGACGATAGTTACCGGATAAGGCGCAGCGGTCGGGCTGAACGGGGGGTTCGTGCACACAGCCCAGCTTGGAGCGAACGACCTACACCGAACTGAGATACCTACAGCGTGAGCTATGAGAAAGCGCCACGCTTCCCGAAGGGAGAAAGGCGGACAGGTATCCGGTAAGCGGCAGGGTCGGAACAGGAGAGCGCACGAGGGAGCTTCCAGGGGGAAACGCCTGGTATCTTTATAGTCCTGTCGGGTTTCGCCACCTCTGACTTGAGCGTCGATTTTTGTGATGCTCGTCAGGGGGGCGGAGCCTATGGAAAAACGCCAGCAACGCGGCCTTTTTACGGTTCCTGGCCTTTTGCTGGCCTTTTGCTCACATGTCCTGCAGGCAG |

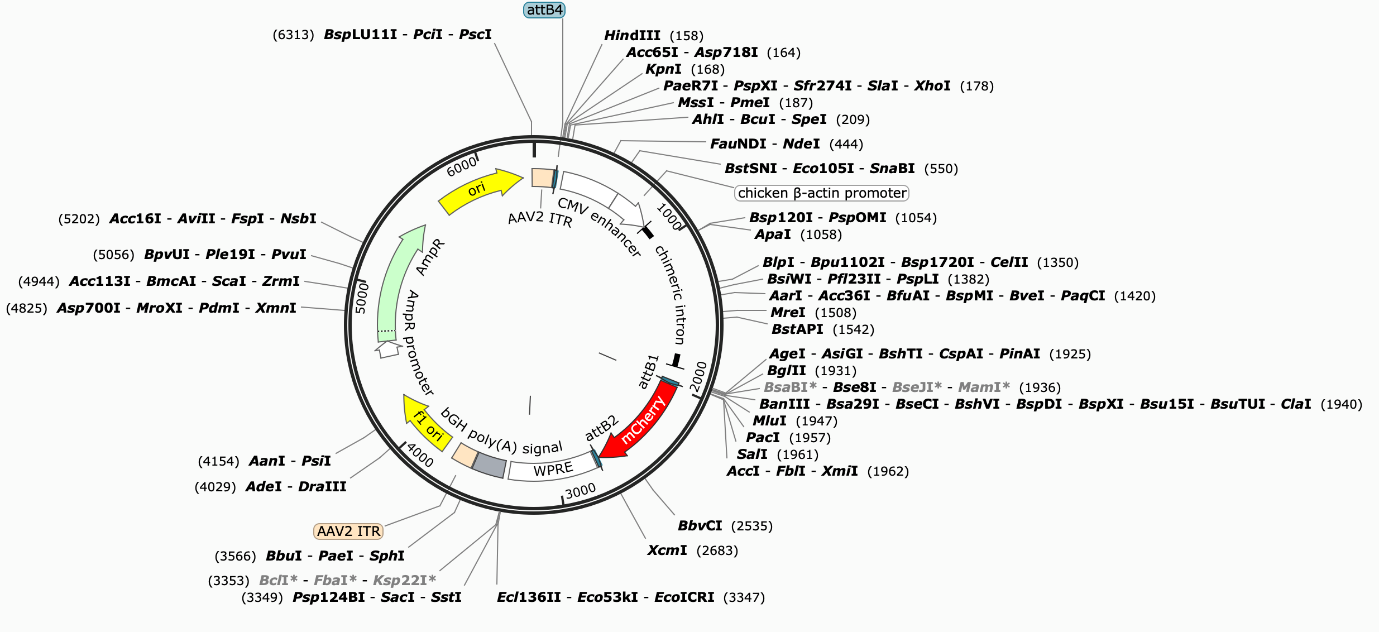
**Plasmid map pDS-000 (pAAV-CAG-mCherry)**

**Plasmid map pDS-001 (pAAV-ACE-tRNA-V3-Stuffer sequence)**

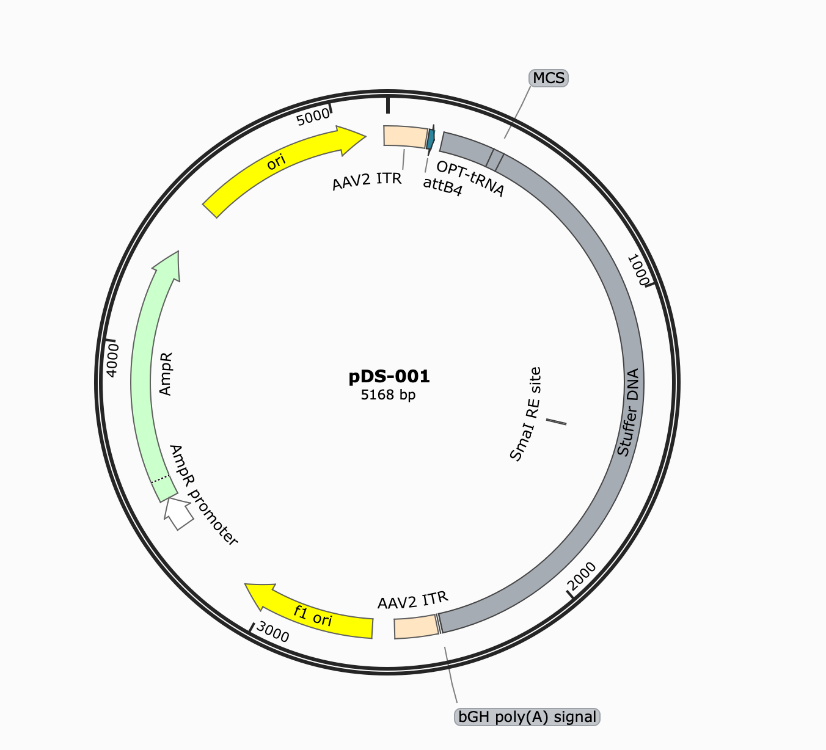

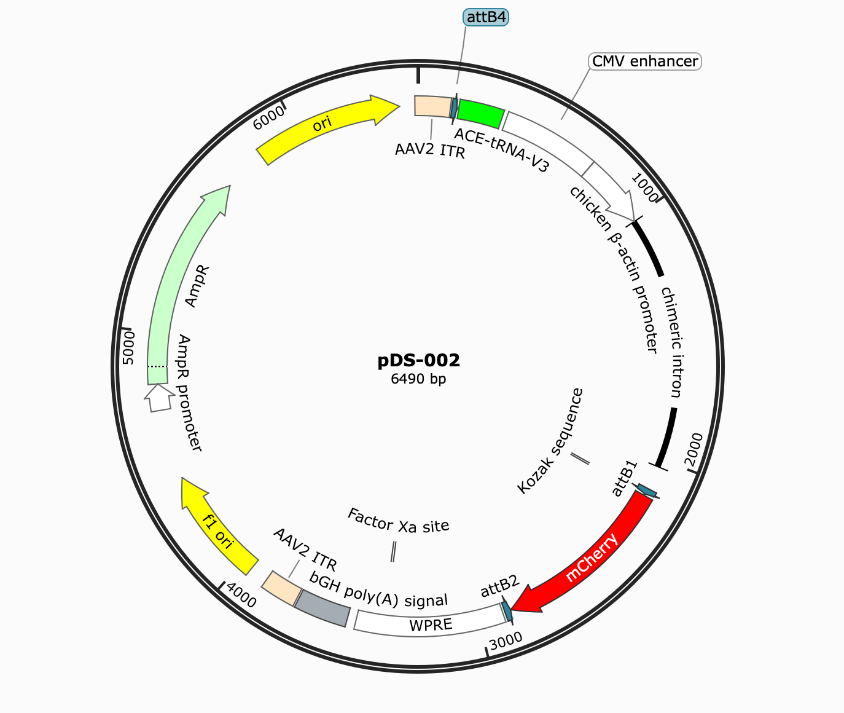
 **Plasmid map pDS-002 (pAAV-ACE-tRNA-V3-CAG-mCherry)**

**Plasmid map pDS-002 (pAAV-ACE-tRNA-V3-CAG-FAM161A-mCherry)**
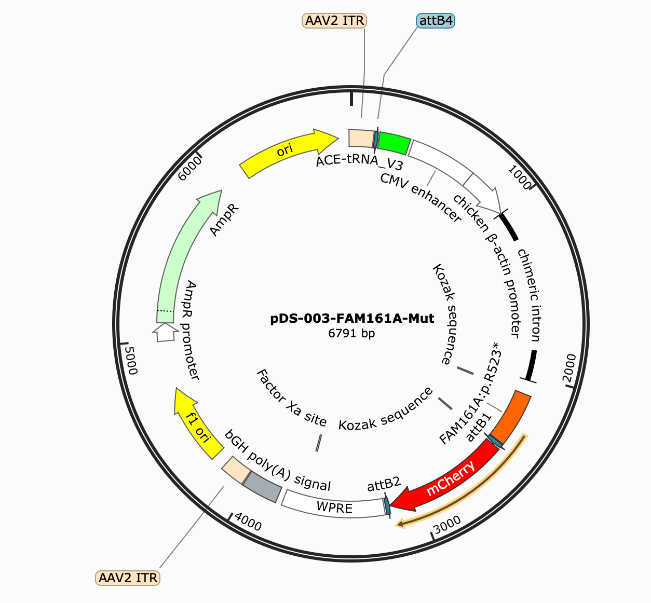

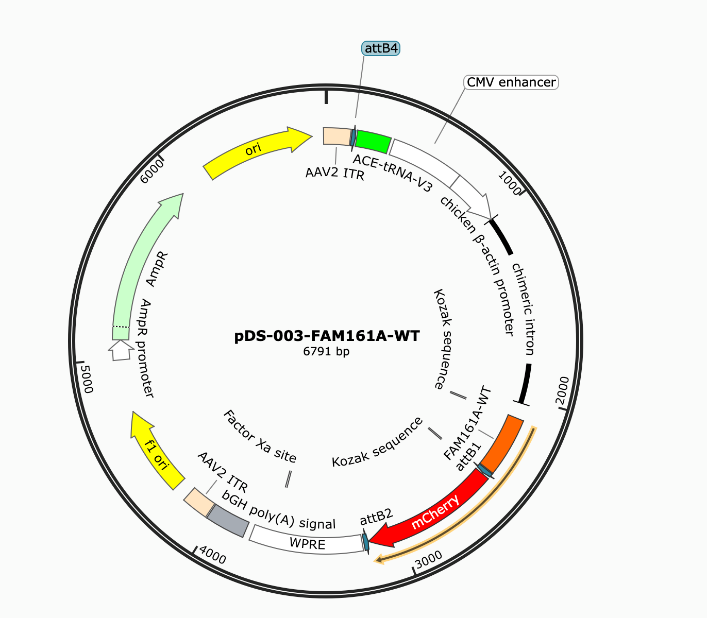

**Supplementary table 5: qPCR Primers used in this study**

| hGAPDH-F | GGAGCGAGATCCCTCCAAAAT |
| --- | --- |
| hGAPDH-R | GGCTGTTGTCATACTTCTCATGG |
| hATF4-F | ATGACCGAAATGAGCTTCCTG |
| hATF4-R | GCTGGAGAACCCATGAGGT |
| hCHOP-F | GGAAACAGAGTGGTCATTCCC |
| hCHOP-R | CTGCTTGAGCCGTTCATTCTC |
| hBIP-F | CATCACGCCGTCCTATGTCG |
| hBIP-R | CGTCAAAGACCGTGTTCTCG |

**Supplementary table 6: list of antibodies used in this study**

| Antibody Name | Host | Company | Catalogue  Number | Application (Dilution) |
| --- | --- | --- | --- | --- |
| GFP | Mouse | Santa Cruz | sc-9996 | 1:1000 (WB) |
| Actin | Mouse | Santa Cruz | c-47778 | 1:200 (WB) |
| GM130 | Mouse | Proeintech | 66662-1-Ig | 1:800 (IF) |
| OCT4 | Mouse | Santa-Cruz | sc-5279 | 1:200 (IF) |
| goat anti-mouse IgG-HRP | Goat | Santa-Cruz | sc-2005 | 1:2500 (WB) |
| mCherry | Rabbit | Sigma-Aldrich | AB356482 | 1:200 (IHC) |
| Cy3 | Cy3-Donkey Anti-Rabbit IgG | Jackson ImmunoResearch Laboratories | 711-165-152 | 1:400 (IHC) |
| Cy3 | Cy3-Donkey Anti-mouse IgG | Jackson ImmunoResearch Laboratories | 715-165-150 | 1:800 (IF) |
